# Impacts of mutation accumulation and order on tumor initiation revealed by engineered murine colorectal cancer organoids

**DOI:** 10.64898/2026.01.05.697837

**Authors:** Yanping Li, Xiaoxin Xie, Daqi Deng, Zhiyuan Sun, Zhaoan Huang, Yisen Tang, Liang Fang, Wei Chen, Qionghua Zhu

## Abstract

Tumorigenesis typically follows a multi-hit trajectory, driven by accumulating oncogenic mutations. Colorectal cancer (CRC) has long served as a paradigmatic model of multi-hit tumorigenesis, characterized by adenoma-carcinoma transition accompanied by acquisition of specific oncogene and tumor suppressor mutations. However, how the temporal order of early mutations influences CRC initiation remains poorly understood. To address this, we established a CRC tumorigenesis model using murine intestinal organoids. By introducing defined combinations of key CRC driver mutations (*Kras*, *Apc*, and *Trp53*) in distinct orders, we systematically investigated how the order of mutation accumulation affects tumor initiation. Our results reveal that the mutation accumulation confers growth advantages in both *in vitro* and *in vivo* models. Strikingly, mutation order also influenced the tumorigenic properties of the organoids. Whereas organoids with *Trp53* loss before or after *Apc* loss similarly affected organoid phenotypes *in vitro* or tumorigenicity in immunodeficient mice, organoids with *Trp53* loss preceding *Apc* inactivation exhibited reduced tumor-forming potential in immunocompetent mice, likely due to their distinct immunological features. Collectively, our study reveals a critical role of ordered mutation accumulation in CRC initiation, an insight that may hold clinical relevance.

## Introduction

Cancer development is an evolutionary process driven by the stepwise accumulation of mutations in key oncogenes and tumor suppressor genes, leading to clonal expansion and tumorigenesis. Early epidemiological studies and population-based statistical analyses have established the paradigm of multistage carcinogenesis and revealed how progressive genetic alterations drive cancer progression^1–3^, especially for retinoblastoma^4^ and colorectal cancer (CRC)^5,6^. Modern multi-omics approaches have further resolved the temporal dynamics of tumor evolution^7–11^, uncovering both conserved and divergent mutational patterns across different malignancies^12–16^. While the minimal number of driver mutations required for tumor initiation has been estimated for several cancer types^5,7,9,17^, inter-tumoral and intra-tumoral heterogeneity reflects varied mutational trajectories during tumorigenesis.

While driver mutations and their approximate timing in tumor evolution are well-characterized, emerging evidence suggests that the temporal order of mutational acquisition, beyond mere presence, can critically affect tumor phenotypes and clinical outcomes. In adrenocortical tumors, for instance, expression of oncogenic Ras^G12V^ followed by dominant-negative p53^DD^ leads to highly malignant, metastatic tumors, whereas the reverse order only leads to benign tumors^18^. This indicates that specific genetic alterations, such as *TP53* deficiency, yield distinct phenotypes depending on their timing during mutagenesis^19^. Similarly, in chronic myeloproliferative neoplasms (MPNs), acquisition of the *JAK2* mutation prior to *TET2* correlates with younger age at diagnosis, polycythemia vera, and a higher risk of thrombosis in patients. In contrast, *TET2*-first patients showed older onset, a lower likelihood of polycythemia vera with essential thrombocythemia, and decreased risk of thrombosis^20,21^. These findings highlighted the importance of systematically investigating the functional impact of mutation order in carcinogenesis^19,22^.

Despite remarkable advances in sequencing technologies and bioinformatics tools which enable comprehensive mutation detection, reconstructing mutational timelines remains challenging. Currently, most of the reconstruction of evolutionary history of individual tumors relies on whole genome/exome sequencing of temporally or spatially separated samples. With the development of computational tools, copy number alterations and single-nucleotide variations can be retrieved from the sequencing data, allowing the inference of partial phylogeny and temporal order of mutations from matched samples and even individual tumors^10,21,23–25^. While clonal and subclonal mutations can be easily timed, the order of truncal mutations that occur early in tumorigenesis is difficult to infer with this strategy. Moreover, most cancers are diagnosed and sampled at advanced symptomatic stages, crucial information regarding premalignant transitions is inevitably lacking, thus limiting the understanding of the order effect on premalignant transitions.

CRC serves as a paradigmatic model for understanding tumor progression since the 1990s, when Fearon and Vogelstein proposed a sequential genetic mutagenesis framework for CRC^6^. Their work revealed that inactivation of tumor suppressors (e.g., *APC*, *TP53*, and *SMAD4*) and activation of oncogenes (e.g., *KRAS*) are important for the neoplastic transition of normal epithelia to malignant tumor cells. Subsequent discoveries of additional driver genes and inter-clonal heterogeneity have further expanded this linear progression model^11,15,26^. Although various theories have been proposed, whether and how the order of mutation acquisition influences tumor outcomes remains elusive.

Conventional CRC models, including cancer cell lines, patient-derived tumor tissues or xenografts (PDXs), and genetically engineered mouse models (GEMMs), are widely used to study tumorigenesis, metastasis, and therapy response. However, these systems lack the capacity to precisely control mutational timing while maintaining isogenic backgrounds. Recent breakthroughs in organoid technology have overcome these limitations by enabling: (i) stepwise genetic manipulation of normal intestinal epithelia, (ii) preservation of *in vivo* physiology with native crypt-villus architecture and lineage hierarchy, and (iii) functional characterization in controlled microenvironments^27,28^.

To systematically investigate how mutation accumulation and order influence CRC initiation, here, we employed CRISPR-engineered intestinal organoids to model the possible tumor evolutionary scenarios. We sequentially introduced three recurrent driver mutations, activation of oncogenic *Kras^G12D^*, and inactivation of *Apc* and *Trp53*, into normal murine intestinal organoids. By doing so, we generated genetically distinct organoid lines with varying numbers, combinations and orders of mutations. Due to the technical challenges associated with sequential genome engineering to create a single-allelic single base mutation and the establishment of fully isogenic organoid clones, our model was initiated with *Kras^G12D^* activation. This strategy provides a stable oncogenic baseline for subsequent mutational engineering and comparative analysis. Although this design does not fully recapitulate the most common evolutionary trajectory of human CRC, where *APC* loss typically occurs first, it provides a controlled and tractable framework for interrogating the phenotypic consequences of mutation order within a defined oncogenic context. For triple-hit organoids, we inverted the acquisition order of *Apc* and *Trp53* mutations, creating two lines, i.e., KAT (*Kras*→*Apc*→*Trp53*) and KTA (*Kras*→*Trp53*→*Apc*). With these engineered organoid lines, we performed comprehensive histopathological and transcriptomic characterization, as well as quantitative assessments of organoid growth *in vitro* and tumor formation *in vivo* using both immunodeficient and immunocompetent hosts. These analyses revealed that mutation accumulation is a key determinant of tumor initiation efficiency. When analyzed individually, *Kras* and *Apc* mutations conferred modest growth advantages, whereas *Trp53* deficiency exhibited a stronger potential for malignant transformation. Notably, although the timing of *Trp53* inactivation relative to *Apc* loss had no impact on organoid growth *in vitro* or tumorigenicity in immunodeficient mice, it substantially affected tumorigenesis in immunocompetent mice: organoids with loss of *Trp53* preceding *Apc* exhibited reduced tumor-forming capacity, likely due to altered tumor-immune interactions. Interestingly, comparative analysis of clinical datasets revealed that engineered KAT organoids, but not KTA, molecularly resembled immune-related phenotypes correlated with poor responses to immune checkpoint therapies in the patients carrying *APC*, *TP53* and *KRAS* mutations. This suggests that the order-dependent effects observed in the engineered murine organoids may also occur in clinical samples, highlighting their potential relevance for guiding the design of therapeutic strategies.

## Results

### Modeling CRC with varied mutation number, combination, and order in murine intestinal organoids

To model the development of early CRC, we selected three of the recurrent driver mutations including *Apc* and *Trp53* loss-of-function mutations and *Kras^G12D^* activation for stepwise mutagenesis. Due to the low-efficiency in generating a *Kras^G12D^*mutation using CRISPR-based knock-in strategy, we chose to employ a transgenic mouse strain carrying Loxp-Stop-Loxp (LSL) alleles of *Kras^G12D^* and *Cas9-tdTomato* (*Kras^LSL-G12D^*^/+^; *Rosa26^CAG-LSL-Cas^*^9^*^-tdTomato/+^*, hereafter designated as SKC for Stop-*Kras^G12D^*-*Cas9*), to establish primary intestinal organoids as previously described (Fig. 1A)^29^. Subsequent infection of adeno-associated virus serotype 9 expressing Cre recombinase (AAV9-Cre) simultaneously activated the expression of *Kras^G12D^*, *Cas9,* and tdTomato, enabling fluorescence-activated cell sorting (FACS) to enrich successfully recombined organoids (designated as K for *Kras^G12D^* activation) (Figs. 1A and S1A). While KRAS mutation may not be the most frequent initiation events, it provided a robust, proliferative baseline that greatly accelerated the generation of a subsequent well-controlled isogenic clone panel. Based on this, we then sequentially introduced mutations in CRC-associated tumor suppressor genes into K organoids^30,31^. For the first-round mutagenesis, we used sgRNAs targeting *Apc* (sgApc-GFP) or *Trp53* (sgTrp53-BFP) to generate two distinct lineages: *Kras^G12D^*; *Apc-KO* (referred to as KA) and *Kras^G12D^; Trp53-KO* (referred to as KT) organoids (Fig. S1B and Table S1). For the second-round mutagenesis, we introduced *Trp53* mutation into KA organoids to generate *Kras^G12D^; Apc-KO; Trp53-KO* (referred to as KAT), and *Apc* mutation into KT organoids to generate *Kras^G12D^; Trp53-KO; Apc-KO* (referred to as KTA), respectively. For each genotype, isogenic organoid clones were isolated by FACS, and targeted mutations were validated through DNA gel electrophoresis, Sanger sequencing, and RT-qPCR accordingly (Figs. 1B, S1A, B and Table S1). To confirm pathway-specific perturbations, we measured the RNA expression of downstream targets of the P53 pathway (*Cdkn1a,* encoding P21) and the Wnt/β-catenin pathway (*Axin2*), both of which showed expected changes in abundance (Fig. 1C). Collectively, we established a comprehensive panel of mouse intestinal organoid lines, including single- (K), double- (KA and KT), and triple-mutant (KAT and KTA) variants, to dissect the dynamics of CRC initiation.

**Figure 1.**
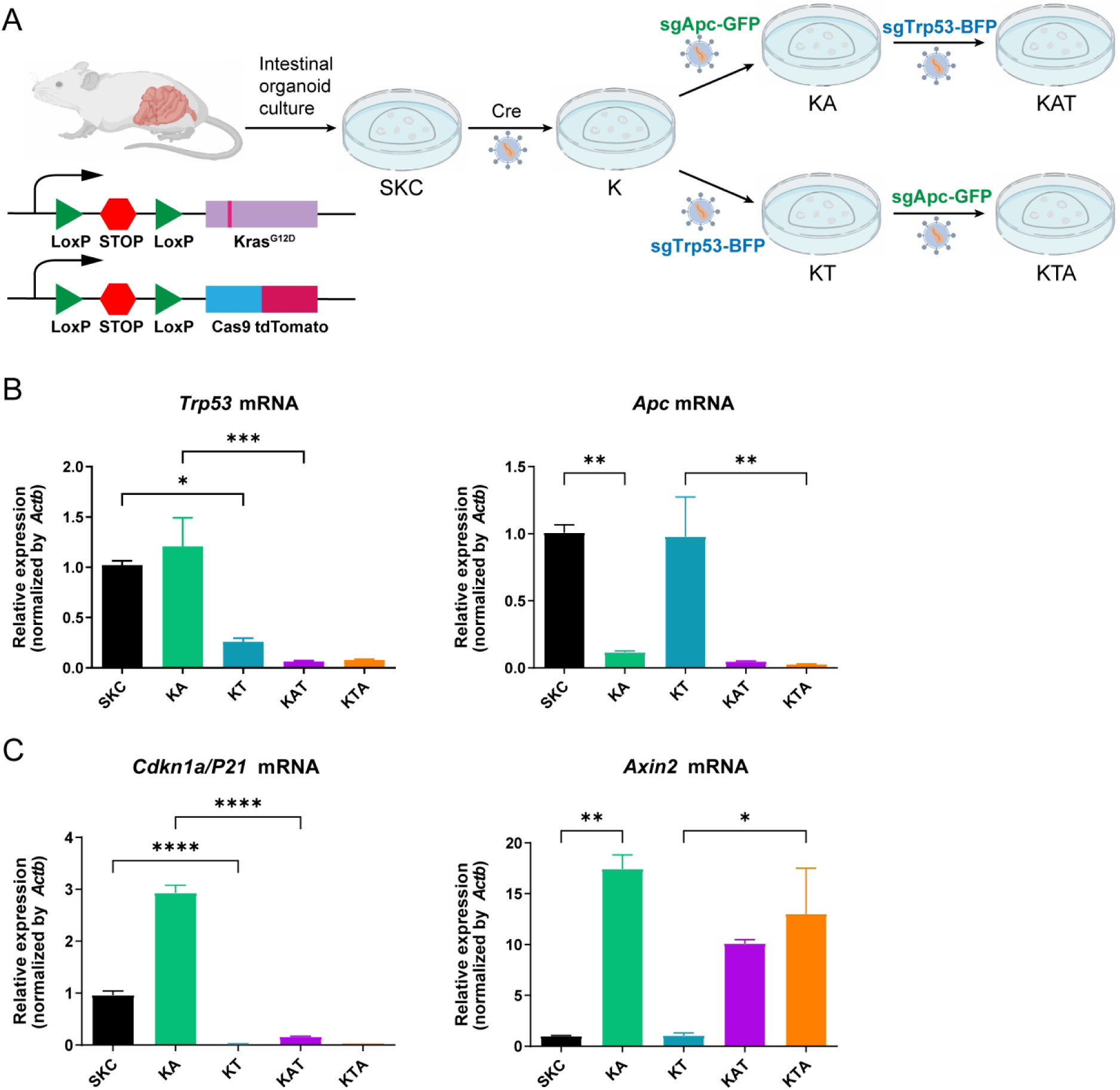
Introduction of CRC driver mutations in murine intestinal organoids. A. Experimental design for sequentially introducing CRC driver mutations into the genomes of primary murine intestinal organoids with Cre recombination and CRISPR-Cas9. Cre recombinase and specific gRNAs were introduced via AAV (adeno-associated virus). K, *Kras^G12D^* activation; A, *Apc* loss; T, *Trp53* loss. B. RT-qPCR analysis of *Trp53* and *Apc* mRNAs expression levels in SKC, KA, KT, KAT and KTA organoid lines. Data is shown as mean ± SEM. (n = 3 biologically independent experiments). C. RT-qPCR analysis of *Cdkn1a* and *Axin2* mRNAs (downstream targets of *Trp53* and *Apc*, respectively) in SKC, KA, KT, KAT and KTA organoid lines. Data is shown as mean ± SEM. (n = 3 biologically independent experiments). Statistical analysis of RT-qPCR was done by One-way ANOVA, adjusted P values: *, P < 0.05; **, P < 0.01; ***, P < 0.001; ****, P < 0.0001. The clones used in this analysis and shown in figure were listed in Table S5.

### Accumulation of mutations drives histopathological progression in engineered organoids

Next, we set out to investigate growth characteristics of engineered organoids. As shown in Figure 2A, all organoid lines predominantly exhibited stem cell-enriched spheroid morphology. Compared to SKC, K and KA organoids, KT and the triple-mutant lines (KAT and KTA) exhibited accelerated expansion and formed larger structures (Figs. 2A, B). These expanding cystic structures could be maintained for 5-7 days per passage. To delve deeper into the cytological impact of distinct CRC mutations, we conducted histological examinations. Hematoxylin and eosin (H&E) staining revealed that engineered organoids formed three-dimensional (3D) structures in Matrigel with epithelial architecture (Fig. 2C). Specifically, SKC and K organoids retained normal monolayered epithelia, whereas KA organoids exhibited early dysplastic features, including epithelium thickening and moderate nuclear enlargement (Figs. 2C, D). Notably, KT and triple-mutant organoids exhibited more severe histologic abnormalities, such as multilayered and disorganized epithelia, focal dysplasia, thinner epithelial regions, and cells with highly pleomorphic and crowded nuclei, showing features of invasiveness. These phenotypes closely resemble the transformation features previously reported in the triple-mutant human colon organoids (*SMAD4-WT; KRAS^G12D^; P53-KO; APC-KO*)^31,32^, underscoring a correlation between mutation load and neoplastic progression in small intestinal organoids.

**Figure 2.**
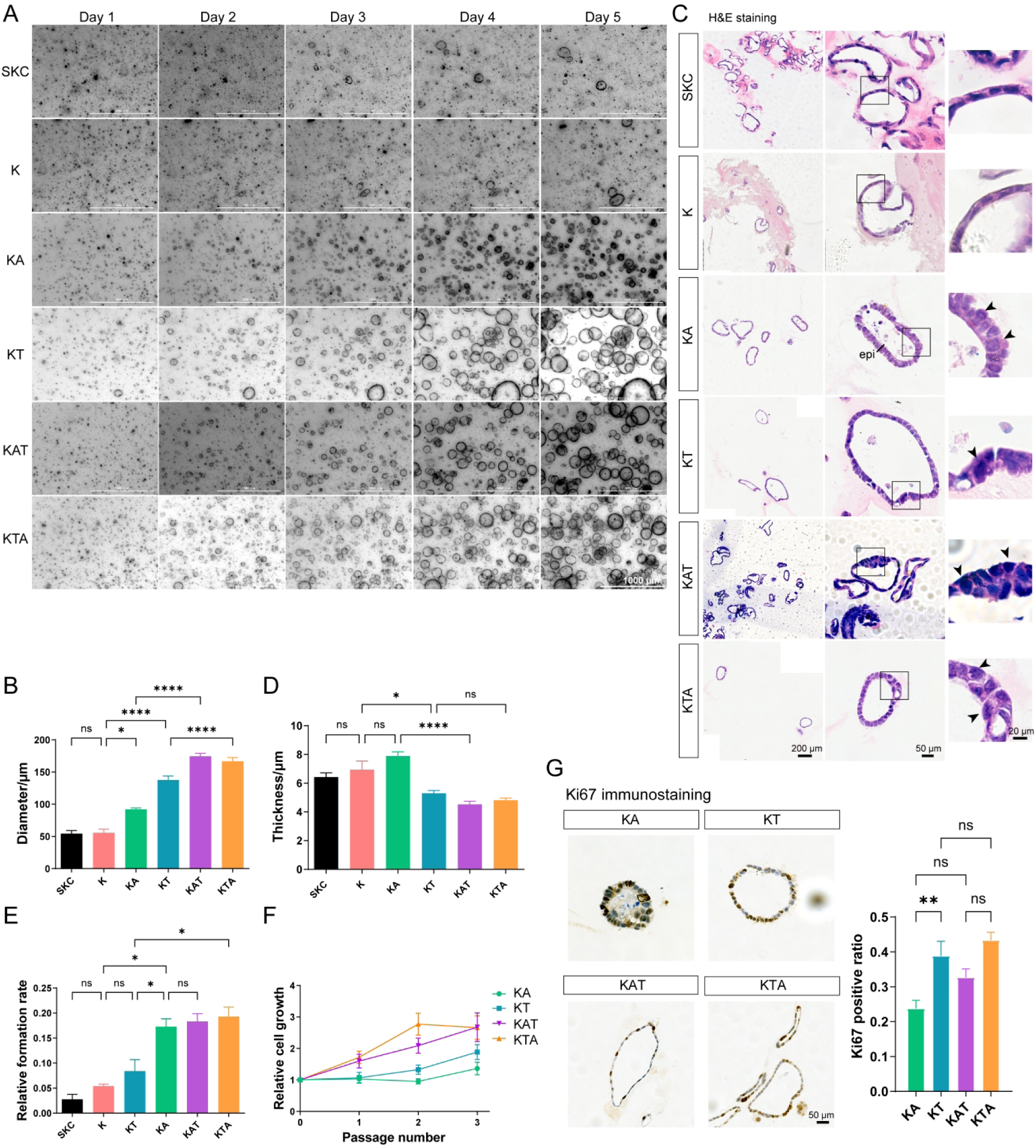
Mutation accumulation drives histopathological progression in engineered organoids. A. Representative bright field images of different organoid lines showing the growth during five consecutive days. Scale bar, 1000 μm. B. Comparison of organoid size at day 5 after single cell seeding. The analysis was performed with N = 2 clonal organoid lines except for SKC and K organoids, and n >= 18 organoids per genotype were measured. C. Histological analysis of different organoid lines by H&E staining, epithelial thickening in KA organoids is indicated by the black bar (epi) and focal dysplasia in KT, nuclear enlargement and disorganized epithelial cells in KA, KT, KAT and KTA organoids is shown by black arrowheads. Scale bar, 200 μm (left panel, overview), 50 μm (middle panel, individual organoid) and 20 μm (right panel, magnification). D. Comparison of organoid epithelium thickness. The analysis was performed with N = 2 clonal organoid lines except for SKC and K, and n >= 6 organoids per genotype were measured. E. Comparison of organoid formation rate at day 5 after single cell seeding. The analysis was performed with N = 2 clonal organoid lines except for SKC and K, and n >= 2 independent formation assays were performed. F. Growth curve of different organoid lines. The analysis was performed with N = 2 clonal organoid lines except for SKC and n >= 3 independent replicates. G. Ki67 immunostaining and proliferative index of mutant organoids. Scale bar, 50 μm. The analysis was performed with N = 2 clonal organoid lines and n = 9 distinct views. Statistical analysis of B, D, E and G were done by One-way ANOVA, P values: ns, non-significant, P > 0.05; *, P < 0.05; **, P < 0.01; ****, P < 0.0001. The clones used in these analyses and shown in figure were listed in Table S5.

To assess the stem cell functionality and proliferative potential of the engineered organoid lines, we quantified their organoid formation efficiency and growth kinetics *in vitro*. Both of SKC and K organoids exhibited low formation efficiency, with K showing a slightly higher, but statistically insignificant, rate than SKC (5% vs. 2%; Figs. 2A, E). Compared with SKC and K organoids, KA organoids displayed a markedly increase in formation efficiency (16%), without a significant increase in growth rate (Figs. 2A, E). In contrast, KT organoids showed enhanced growth over K and KA organoids (Fig. 2A), along with a higher organoid formation efficiency compared to K organoids, though this difference was not statistically significant (Fig. 2E). As expected, triple-mutant lines (KAT and KTA) exhibited the highest clonogenic capacity (17-18%) and accelerated proliferation. Consistently, Ki67 immunostaining demonstrated elevated proliferative activity in all *Trp53*-deficient lines. Specifically, both KT and KTA organoids showed a statistically significant increase in proliferation compared to KA controls, while KAT organoids showed a strong, albeit statistically insignificant, trend toward higher rate (Figs. 2A, E-G). This overall enhancement of proliferation in p53-null backgrounds is in line with the known role of Trp53 in cell cycle regulation^32^. As for the triple-mutant organoids with reverse orders, they displayed similar histomorphological features, as well as comparable clonogenic and proliferative capacities, resembling previously reported models generated either via sequential mutation (A → K → T) or simultaneous induction of all three mutations^33,34^. Taken together, these results suggest that mutation burden correlates with both progressive morphological and functional transformation. Notably, *Trp53* inactivation exerts a dominant effect in promoting proliferation, while *Apc* loss contributes to enhanced organoid-forming capacity.

### Mutation accumulation determines the tumorigenicity of engineered organoids

To investigate how the accumulation of driver mutations influences tumorigenesis, we performed subcutaneous transplantation of dissociated cells from single- (K), double- (KA, KT), and triple-mutant (KAT, KTA) organoids into immunodeficient B-NDG (NSG; NOD-*Prkdc^scid^IL2rg^tm^*^1^) mice for up to three months. In this transplantation experiments, due to the varied tumorigenicity of the engineered organoids, the tumors were harvested at different time points once they met the collection criteria (see Methods) to examine their features. As anticipated, single-mutant K organoids failed to form detectable nodules, while double-mutant KA organoids only gave rise to small and soft plugs (Figs. 3A, B). Notably, KT organoids generated measurable solid tumors with significantly greater weight and volume compared to KA-derived lesions (Figs. 3A, B, S2A and B), demonstrating stronger oncogenic synergy between *Trp53* deficiency *and Kras^G12D^* activation than between *Apc* loss and *Kras^G12D^* activation. Strikingly, triple-mutant organoids (KTA and KAT) exhibited the most robust tumorigenicity, forming visible and occasionally voluminous tumors (Figs. 3A, B, S2A and B). These results demonstrate that tumor latency inversely correlates with mutation burden, consistent with previous mutagenesis studies^34,35^. Together, our findings indicate that combined *Kras^G12D^* activation and *Trp53* loss is necessary for tumor initiation, and the capacity for malignant tumor formation ability increases proportionally with mutational load.

**Figure 3.**
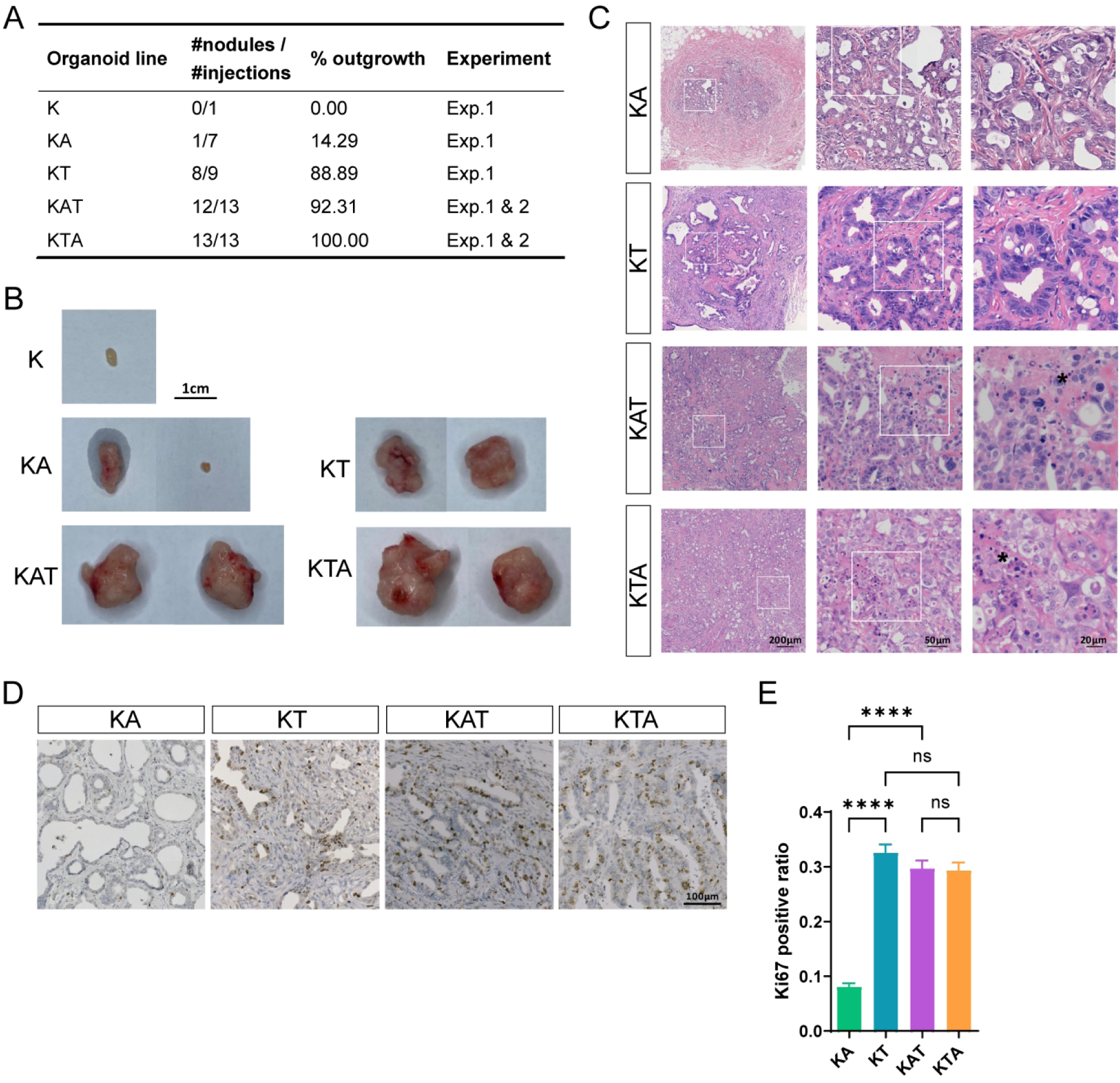
Mutation burden-dependent tumorigenesis of engineered organoids. A. Tumor outgrowth of organoid lines with different mutations in immunodeficient mice, showing the combined data from two independent experiments (see Methods for criteria defining outgrowth). B. Representative images of subcutaneous tumors developed from injected organoids with different mutations. All tumors were collected upon termination of the first experiment, with varying endpoint times. Scale bar, 1 cm. C. Representative H&E staining images showing the tumor morphology of KA, KT, KAT, and KTA tumors, necrosis was indicated with black asterisks. Scale bar, 200 μm (left panel, overview), 50 μm (middel panel), and 20 μm (right panel) for magnification. D. Representative images of Ki67 immunostaining showing proliferating cells in KA, KT, KAT, and KTA tumors from the first experiment. Scale bar, 100 μm. E. Ki67 index in indicated tumors. The analysis was performed with N >= 2 clonal organoid lines and n = 9 distinct views, statistical analysis was done by One-way ANOVA, adjusted P values: ns, non-significant, P > 0.05; ****, P < 0.0001. The clones used in these analyses and shown in figure were listed in Table S5.

Histologically, H&E analysis revealed mutation-associated tumor architectures (Fig. 3C). Both KA- and KT-derived tumors maintained organized and moderately differentiated epithelial structures with stromal encapsulation, resembling early-stage colorectal adenoma. Immunohistochemical analysis showed significantly higher Ki67 labeling index in *Trp53*-deficient (KT, KAT and KTA) tumors compared to KA lesions (Figs. 3D, E), consistent with their enhanced proliferation capacity observed *in vitro*. In contrast, triple-mutant tumors displayed poorly differentiated adenocarcinomas, with irregular and compacted glandular structures and extensive necrosis, a pattern consistent with rapid tumor expansion (Figs. S2A, B), as also seen in orthotopic transplants^32^.

Next, we asked whether the order of mutation acquisition influences tumor phenotypes. Subcutaneous transplantation to immunodeficient mice demonstrated equivalent tumorigenic potential between KAT and KTA. Based on our predefined outgrowth criteria (see Methods), there were no significant differences in tumor incidence, latency, or final tumor mass (the first experiment) (Figs. 3A-C and S2B). To further explore this aspect, we repeated the experiment for the KAT and KTA lines and strictly controlled the tumor formation within the same period in the second experiment, but again found no significant differences (Figs. S2C, D, see Methods). Histopathological evaluation revealed that KAT tumors had a slightly higher proportion of tumor cells, as shown by epithelial cell marker keratin 20 (KRT20) immunostaining (Fig. S2E), suggesting enhanced colonization and outgrowth potential. However, no significant differences in Ki67 index were observed between KAT and KTA tumor cells (Figs. 3E).

Taken together, these transplantation assays demonstrate a clear mutation burden-dependent trend in tumorigenesis. *Apc* loss initiates transformation but requires cooperative genetic alterations for malignant transformation^36^, whereas *Trp53* ablation promotes proliferative expansion and pathophysiological progression. This underscores their non-redundant and synergistic roles in tumor development. Although mutation order did not affect overall tumorigenicity in immunodeficient hosts, the enhanced epithelial colonization (KRT20^+^ fraction) in KAT tumors suggests that mutation order may subtly influence tumor cell fitness, independent of proliferation rate.

### Sequential mutational acquisition drives dynamic transcriptional changes

To delineate the molecular consequences of stepwise mutagenesis, we conducted transcriptomic analysis across all engineered organoid lines. Principal component analysis (PCA) revealed that the trajectory of transcriptional changes closely mirrored the stepwise acquisition of genetic manipulations, with principal component 1 (PC1) capturing gene expression changes associated with mutation acquisition, and PC2 showing two evolutionary paths corresponding to distinct mutation orders (Fig. 4A). As expected, the number of differentially expressed genes (DEGs) relative to SKC controls gradually increased with accumulation of mutations (Fig. 4B). Gene set enrichment analysis (GSEA) further validated the transcriptomic effect of gene-specific perturbation: *Kras^G12D^* activation resulted in upregulation of KRAS signaling, loss of *Trp53* caused profound suppression of P53 pathway, and *Apc* inactivation triggered the activation of Wnt/β-catenin signaling (Figs. S3A, B).

**Figure 4.**
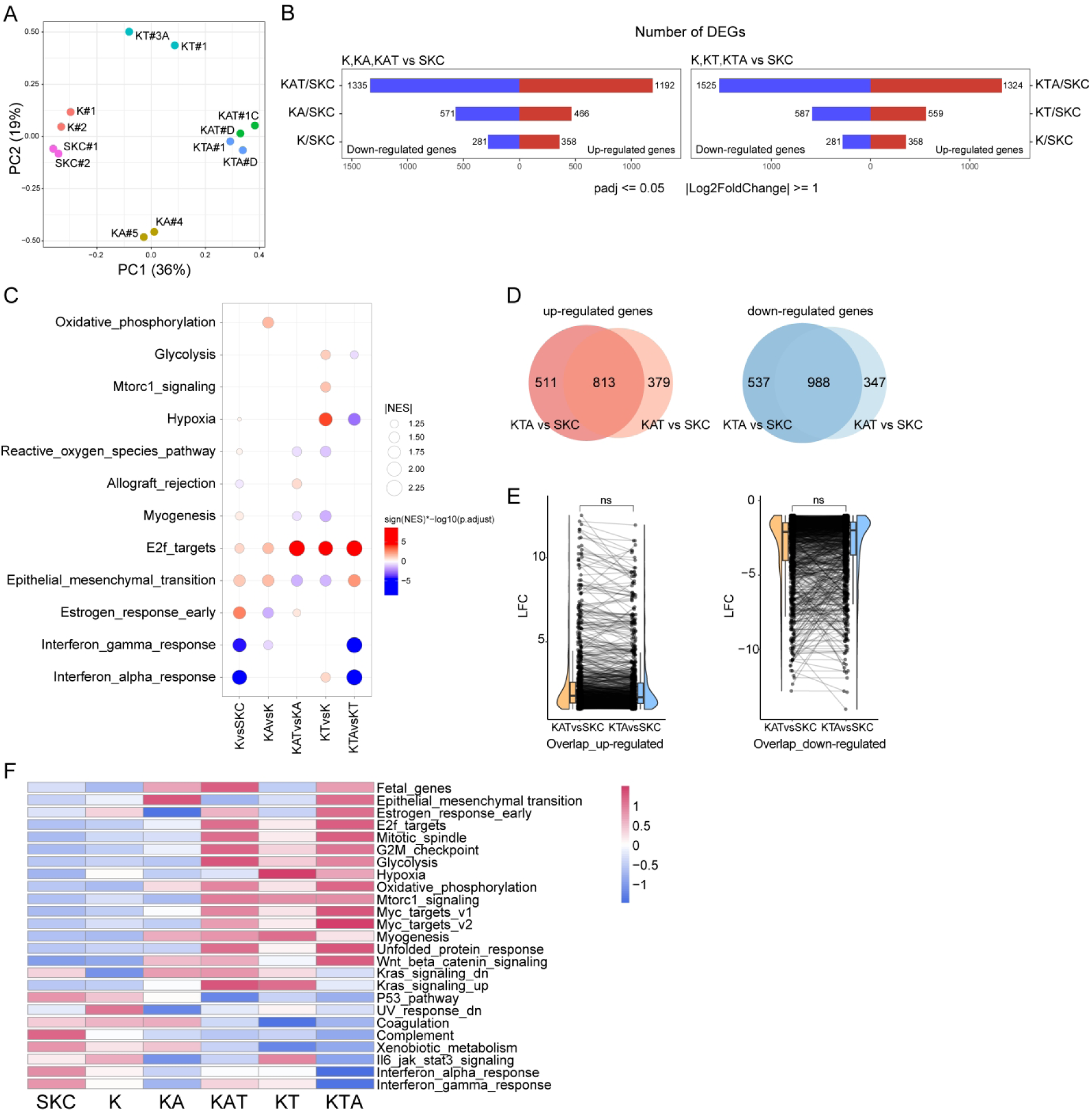
Transcriptional changes during stepwise tumorigenesis. A. Principal component analysis (PCA) of transcriptomic profiles from different organoid lines, and organoids with different genotypes are labeled by colors. PC1 and PC2 explain 36% and 19% of the total variance, respectively. B. Bar plot showing the number of differentially expressed genes (DEGs) along the mutagenesis, all comparisons were performed relative to SKC organoids. The DEGs were defined as genes with Log2 Fold Change (LFC) >= 1, adjusted P values (padj) <= 0.05. Red bars represent up-regulated genes, and blue bars represent down-regulated genes. C. GSEA summary showing the key biological pathway dysregulations upon *Kras^G12D^* mutation and subsequent *Apc* or *Trp53* loss. Red and blue dots represent activation and suppression of indicated pathways, respectively. The color is scaled as indicated by adjusted P values. D. Venn plots showing the overlap of DEGs from KAT vs. SKC and KTA vs. SKC comparisons. Up-and down-regulated genes are colored as red and blue. E. Raincloud plots showing the comparable magnitudes in expression changes (indicated by paired lines connecting two points in two groups) of the overlapped 813 up-regulated and 988 down-regulated genes, suggesting mutation order minimally impacts overall transcriptional output, significant analysis was done by Wilcoxon test, ns: P > 0.05. F. Summary dynamics of transcriptional changes in selected gene sets, as indicated. The bar corresponding to each stage is colored by the log transformed gene expression z-score of each gene set. The clones used in transcriptional analyses were listed in Table S5.

The initial single *Kras^G12D^* oncogenic mutation induced a preneoplastic transcriptional state characterized by activation of reactive oxygen species pathway, myogenesis, E2f targets, epithelial-mesenchymal transition, and estrogen response early, alongside attenuated interferon responses (Fig. 4C). These changes may collectively contribute to the cell proliferation and invasion^37^. Introduction of *Apc* or *Trp53* mutation in K organoids resulted in enhanced activation of E2f targets (normalized enrichment scores, NES= 1.542 in KA; NES= 2.057 in KT), potentially contributing to tumor cell proliferation as described earlier. Compared to KA organoids, which were significantly activated in oxidative phosphorylation, KT organoids preferentially activated glycolysis, mTORC1 signaling, and hypoxia pathways. In addition, consistent with their distinct morphological (Fig. 2) and tumorigenic phenotypes (Fig. 3), KA and KT organoids showed significant transcriptional divergence, with 623 genes upregulated and 639 downregulated between the two genotypes (Fig. S3C). Gene Ontology (GO) analysis uncovered KA-specific up-regulation of cell adhesion and down-regulation of cell differentiation regulators and inflammatory responses (Fig. S3D). This transcriptional pattern likely accounts for the epithelial thickening observed in KA organoids, as enhanced cell adhesion combined with impaired epithelial differentiation may underlie this morphological change. These observations align with known roles of APC inactivation in regulating epithelial polarity and differentiation^36,38^.

In organoids harboring all three mutations (*Kras, Apc,* and *Trp53*), gene expression changes largely overlapped between the two triple-mutant genotypes (KAT and KTA) (Fig. 4D), with no significant differences in commonly altered genes relative to SKC controls (Fig.4 E). Both triple-mutant organoids exhibited significant CRC-intrinsic patterns, including Wnt/β-catenin signaling activation, P53 pathway suppression and further enhanced activation of E2f targets (Fig. 4F), reflecting synergistic alterations in core oncogenic pathways. Besides, the triple-hit organoids recapitulated fetal-like transcriptional signatures as previously described in CRCs^39,40^, and showed hallmark features of malignant transformation, including increased expression of cell replication-related genes, alterations in oxidative metabolism, and a progressive erosion of interferon responses pathways following *Apc* loss^41^. Together, these findings underscore that transcriptomic alterations were directly modulated by the stepwise introduction of driver mutations.

Next, we investigated whether early mutation-induced transcriptional changes were preserved upon additional mutagenesis. To assess the temporal dynamics of these changes, we compared the gene expression profiles across successive mutational stages (all relative to SKC; Fig. S4A). Most transcriptional alterations observed in K organoids were not fully maintained in subsequent stages (KA and KT). Specifically, only 8.4% (30 out of 358) up-regulated and 28.5% (80 out of 281) down-regulated genes in K vs. SKC overlapped with the DEGs in KA vs. SKC (marked as “d” and “g” in Fig. S4B and Table S2). Likewise, 21.2% (76 out of 358) up-regulated and 32.7% (92 out of 281) down-regulated genes in K vs. SKC showed consistent changes in KT vs. SKC (marked as “g” and “e” in Fig. S4C and Table S2). This result suggested that early transcriptional responses were transient and likely represent an adaptive state insufficient for sustained transformation. In contrast, the transcriptional programs established at KA and KT stages remained relatively stable following further mutagenesis. For instance, 56.2% (262 out of 466) up-regulated and 60.2% (344 out of 571) down-regulated genes in KA vs. SKC were conserved in KAT vs. SKC (marked as “g” and “e” in Fig. S4B and Table S2). Similarly, 50% (280 out of 559) up-regulated and 72.9% (428 out of 587) down-regulated genes in KT vs. SKC remained consistently dysregulated in KTA vs. SKC (marked as “g” and “e” in Fig. S4C and Table S2). This stability was further supported by higher correlations in gene expression fold-changes between KA and KAT, and between KT and KTA, respectively (Fig. S4A). Collectively, our data demonstrate that the accumulation of CRC driver mutations governs the temporal evolution of transcriptional programs, with early-stage changes being transient and unstable, whereas later-stage changes gaining increasing stability.

### Mutation order influences transcriptional profiles through context-specific effects of mutations

Transcriptomic analysis of triple-mutant organoids (KAT vs. KTA) identified 297 DEGs (179 up- and 118 down-regulated genes) associated with mutation order (Fig. 5A). Notably, KAT organoids exhibited upregulation of immune response genes, alongside downregulation of cell adhesion and lipid metabolism pathways (Fig. 5B). We speculate that the order of mutation acquisition may differentially shape transcriptional programs by modulating the functional impact of *Apc* and/or *Trp53* loss within specific genetic contexts.

**Figure 5.**
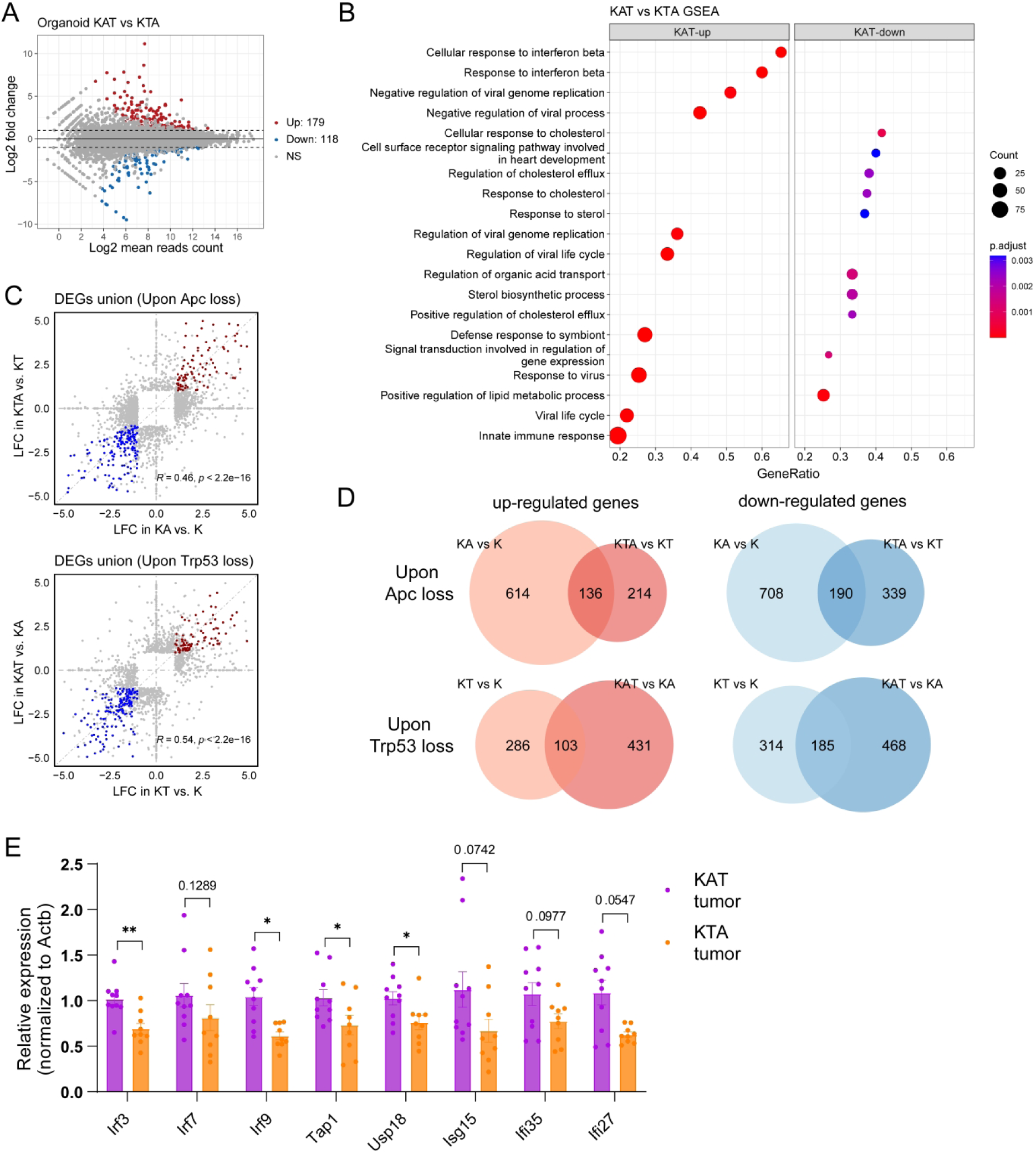
Comparative transcriptomic analysis revealing conserved and context-specific gene expression changes induced by the same genetic mutation. A. MA-plot showing the DEGs between KAT and KTA organoids, with 179 up-regulated genes and 118 down-regulated genes, defined as genes with Log2 Fold Change (LFC) >= 1, adjusted P values (padj) <= 0.05. B. GSEA enrichment showing the transcriptomic differences between KAT and KTA organoids, with interferon responses being most significantly activated and lipid metabolism being suppressed in KAT organoids compared to KTA organoids. C. Correlation of gene expression changes showing the LFC of up- and down-regulated genes (marked as red and blue dots) in indicated transcriptional comparison upon *Apc* (up panel) and *Trp53* (lower panel) loss at different genetic background. D. Venn plots showing the overlap of the consistent up- (left panel) and down- (right panel) regulated genes when acquiring *Apc* or *Trp53* mutation at different genetic background. E. RT-qPCR analysis of immune-related genes in KAT and KTA tumors derived from immunodeficient mice, and data is shown as mean and SEM. (n = 10 KAT tumors and n = 9 KTA tumors). Significance analysis was done by Wilcoxon test, P values are indicated in non-significant groups; *, P < 0.05; **, P < 0.01. The clones used in these analyses were listed in Table S5.

To further explore this, we examined the transcriptional consequences of *Apc* loss in distinct mutational backgrounds. While global transcriptional changes were broadly similar (Fig. 5C), for instance, Wnt signaling activation was a consistent feature of *Apc* loss regardless of genetic background (136 shared upregulated genes, Fig. S5A), substantial context-dependent differences emerged (Fig. 5D). Specifically, early *Apc* loss (K→KA) induced a broader transcriptional shift (750 upregulated and 898 downregulated genes) than late *Apc* loss (KT→KTA: 350 upregulated and 529 downregulated genes; Figs. S3A and 5D). The *Apc* loss preferentially upregulated genes involved in nuclear division and chromosome segregation, while downregulating cytoskeletal organization and differentiation pathways (Fig. S5C), consistent with known roles of APC in regulating DNA repair and cell cycle progression^42^. In contrast, late *Apc* inactivation primarily activated canonical and non-canonical Wnt signaling without inducing additional proliferative programs, while downregulated genes were predominantly related to innate immune response (Fig. S5D).

A similar pattern of conserved and context-dependent responses was observed for *Trp53* inactivation. While certain canonical changes, such as increased expression of nuclear division genes and suppression of apoptotic pathways, were consistently observed regardless of genetic background (Fig. S5B), certain pathways were differentially regulated in a time-specific manner. Specifically, late *Trp53* loss (KA→KAT) triggered more DEGs (534 upregulated and 653 downregulated genes) than early loss (K→KT; 389 upregulated and 499 downregulated genes; Figs. 5D and S3A). Early *Trp53* loss selectively downregulated genes associated with oxidative metabolism (response to oxidative stress) and migration-related pathways (wound healing; Fig. S5E), whereas late loss (KA→KAT) upregulated hypoxia response genes (Fig. S5F).

Collectively, these findings confirm the well-established oncogenic roles of *Apc* and *Trp53*, while highlighting the influence of genetic background on the transcriptional reprogramming by the same perturbation during tumorigenesis. Although triple-mutant organoids (KAT and KTA) shared broadly similar gene expression landscapes, they exhibited subtle yet functionally relevant differences in immune modulation, differentiation, and metabolic adaptation. The molecular differences between KAT and KTA organoids likely arose from genetic background-dependent gene regulatory mechanisms and indicate that mutation order can fine-tune tumor-associated transcriptional programs, even in the presence of identical driver mutations.

### Mutation order-dependent tumorigenesis in immunocompetent mice

Accumulating evidence demonstrates that components of the tumor microenvironment, including stromal and immune cells, can profoundly modulate tumor progression^33,43^. Given that KAT and KTA organoids exhibited distinct immune-related transcriptional profiles yet comparable tumorigenic potential in immunodeficient mice, we firstly checked whether the differential expression in genes related to immune response observed between KAT and KTA organoids is maintained *in vivo*. For this purpose, the most significant DEGs including interferon regulatory factors (*Irf3*, *Irf7* and *Irf9*), antigen presenting related genes *Tap1*, as well as IFN-stimulated genes (ISGs) (*Usp18*, *Isg15*, *Ifi27* and *Ifi35*), were selected as candidate genes for RT-qPCR analysis of KAT and KTA tumor samples. As shown in Figure 5E, KAT tumors from immunodeficient mice exhibited significantly higher expression of most immune-related genes compared to KTA tumors. This result suggests that the differential immune responses are cell-intrinsic transcriptomic signatures which are maintained *in vivo*.

Since the immune-related signatures were consistently preserved in tumors from immunodeficient mice, we then investigated how the immune environment affects their tumorigenic behavior in immunocompetent mice. Strikingly, KTA organoids failed to form tumors (0/10 injections), whereas KAT organoids generated detectable tumors in 50% of cases (5/10 injections; Fig. 6A). The complete absence of KTA tumor outgrowth, despite their robust growth in immunodeficient mice, underscores a pivotal role for host immune responses in selectively restricting tumorigenesis. Although KAT organoids successfully formed tumors under immunocompetent conditions, tumor growth was significantly attenuated compared to that in immunodeficient hosts (Fig. 6B, data from the second tumor formation experiment, see Methods), which resulted in smaller final tumor masses even after extended observation (Fig. 6C). These findings demonstrate that immune surveillance provides a stringent barrier for tumor progression. Additionally, only limited infiltration of CD4^+^ and CD8^+^ T cells was observed, primarily localized to the tumor periphery (Fig. 6D), but this limited infiltration of T cells may be sufficient to cause the attenuated tumorigenicity. Compared to KTA, the outgrowth of KAT tumors under immune pressure indicates these cells may possess unique adaptive mechanisms to evade immune clearance and maintain proliferative capacity. Together, these results highlight mutation order as a critical determinant of tumor-immune interactions and underscore its role in shaping evolutionary trajectory of tumorigenesis in immunocompetent hosts.

**Figure 6.**
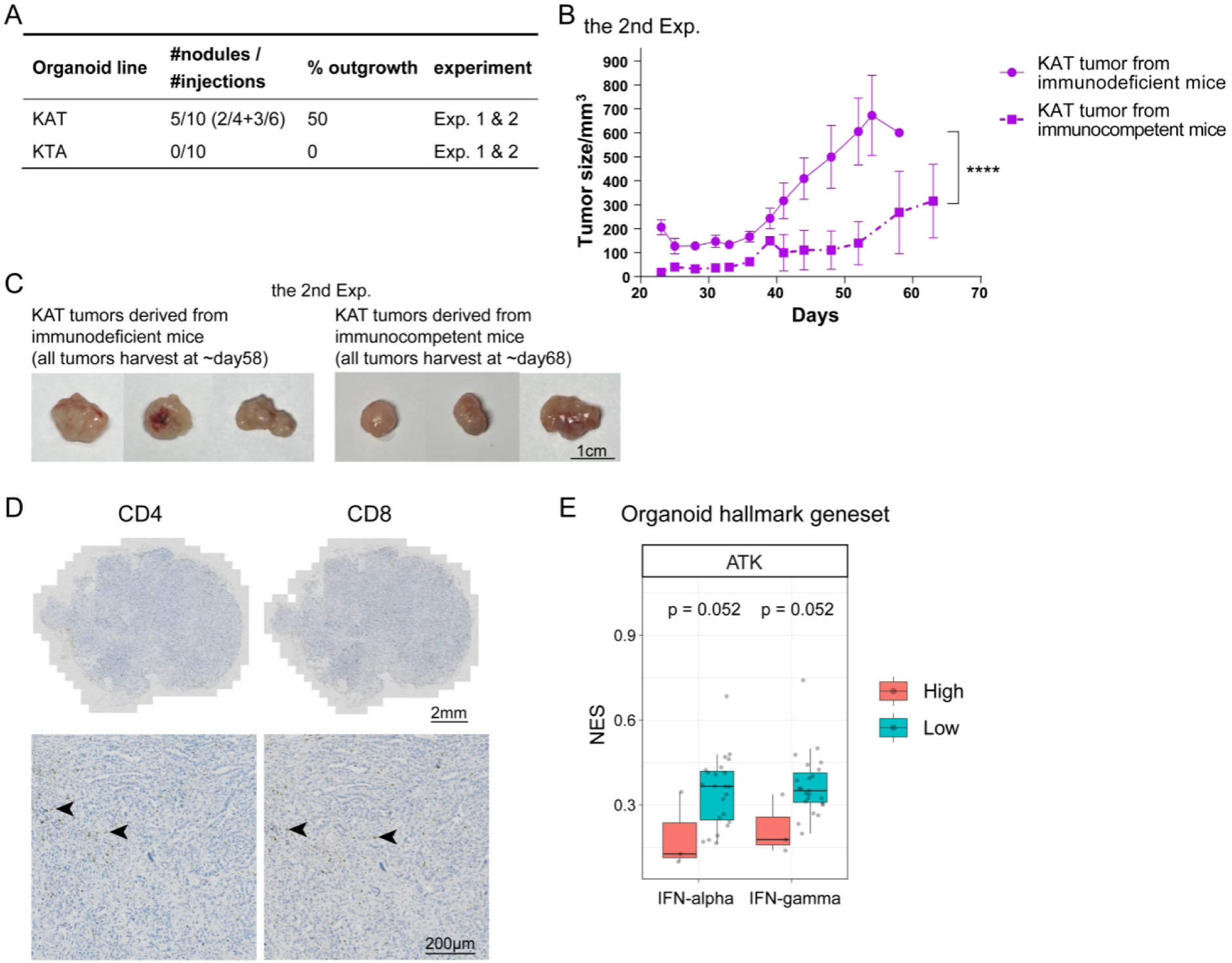
Tumor formation of KAT and KTA organoids in immunocompetent mice and investigation of clinical relevance. A. Tumor outgrowth of KAT and KTA organoids in immunocompetent mice, showing the combined data from two independent experiments. B. Growth kinetics of KAT tumors (of the same isogenic clone) derived from both immunodeficient (n = 3 tumors) and immunocompetent (n = 3 tumors) hosts overtime, data from the second experiment. C. Representative images of subcutaneous KAT tumors (of the same isogenic clone) derived from both immunodeficient (n = 3 tumors, the second experiment) and immunocompetent (n = 3 tumors, the second experiment) hosts at the experimental endpoint. Scale bar, 1 cm. D. Representative images of CD4 and CD8 (T cell) immunostaining with positive cells indicated by black arrows. Scale bar, 2 mm (up panel, overview) and 200 μm (down panel, magnification). E. Bar plot showing the NES distribution of IFN-alpha and IFN-gamma signatures in ATK CRCs, which were generated from transcriptome comparison of KAT and KTA organoids (see Table S2, named organoid hallmark geneset here), red dots represent IFN-high IP tumors (n = 2) and blue dots represent IFN-low IP tumors (n = 12). Statistical analysis was done by Wilcoxon test, and P values are indicated in the figure. The clones used in these analyses were listed in Table S5.

### Clinical Relevance of Mutation Order in CRC Patients

To assess the clinical relevance of our findings on mutation order-dependent effects, we analyzed a multi-regional transcriptomic dataset of CRC patients, which characterized both tumor epithelium (Te) and tumor-associated stroma (Ts) and then categorized tumors into IFN-high and IFN-low immunophenotypes (IPs)^44^. Notably, IFN-high IP tumors showed superior responsiveness to immune checkpoint inhibitors (ICIs) compared to IFN-low IP tumors. Among the 59 CRC tumors analyzed, we identified 14 samples harboring concurrent mutations in *APC*, *TP53,* and *KRAS*, termed ATK CRCs. These ATK tumors exhibited overall lower NES for immune signatures compared to other CRC samples (Table S4). Among the 14 ATK tumors, 12 were classified as IFN-low IP, suggesting that the ATK CRCs preferentially adopt an immune-suppressed tumor microenvironment aligning with their clinical prevalence, aggressive behavior, and poor ICI responses.

To further explore the correspondence between our organoid results and clinical findings, we performed single-sample gene set enrichment analysis (ssGSEA) on the 14 ATK tumors using the IFN response-geneset derived from the of KAT vs. KTA comparison (Table S2). We observed an overall positive association (NES > 0) of ATK CRCs to IFN response signatures (Fig. 6E). Interestingly, the IFN-low IP subgroup tended to show higher NES values for the IFNα and IFNγ responsive signatures that were upregulated in KAT organoids, although the statistical significance was modest (Fig. 6E). This suggests that our KAT organoids partially recapitulate key transcriptional features of the IFN-low IP ATK tumors from CRC patients. This finding points to the temporal order between *APC* and *TP53* as a potential regulator of tumor-immune interactions during CRC development, which may have important implications for CRC prognosis and the rational design of immunotherapeutic strategies.

## Discussion

Using organoids to model intestinal malignancies with sequential acquisition of driver mutations, we systematically investigated phenotypic and transcriptomic characteristics of organoids with varied mutation numbers, combinations, and orders. We observed progressive malignant transformation with increasing mutation burden in our organoid models both *in vitro* and *in vivo*. By systematically dissecting the role of specific mutations, our study enhanced the understanding of both the intrinsic effects of individual mutations and their cumulative, order-dependent impacts on carcinogenesis: *Apc* loss (A) drives dedifferentiation via Wnt activation without inducing full malignant transformation, while *Trp53* loss (T) accelerates tumor progression. Notably, *Kras^G12D^* (K) alone or with *Apc* loss (KA) generated only well-differentiated transplants, whereas KT organoids led to hyperproliferative tumors. The triple-hit KAT/KTA organoids exhibited features of adenocarcinoma, with mixed glandular and poorly differentiated areas, representing the conversion of primary intestinal epithelium to malignancy.

While mutation order is recently hypothesized to influence tumor initiation and progression^19–22,45^, experimental validation has been limited due to the lack of suitable research systems. A recent study demonstrated that *FBXW7*/*APC* mutation order influences clinical phenotypes of CRC based on human colon organoids^45^. Here, by reversing the order of *Trp53* and *Apc* loss in *Kras*-mutant backgrounds (KAT vs. KTA), we uncovered an order-dependent effect specifically in immunocompetent hosts. Interestingly, while growth advantage and tumorigenicity remained unchanged in immunodeficient mice, KAT organoids, but not KTA, formed tumors in immunocompetent mice. This suggests that tumor progression is shaped not only by cell-intrinsic changes but also by interactions with the microenvironment. The elevated immune-response gene expression in KAT organoids and tumors implies order-dependent immune crosstalk, consistent with reports of co-evolution between mutant clones and their environment^46^. Although the transcriptional differences between KAT and KTA organoids were relatively modest under monoculture conditions, these intrinsic differences may become amplified under immune selection pressure *in vivo*. In particular, differential interferon responsiveness, antigen presentation capacity, or tumor cell-intrinsic immune visibility programs may alter susceptibility to immune-mediated clearance^47,48^. Future studies incorporating deeper immune profiling and functional immune assays will be important to clarify the mechanisms underlying these order-dependent phenotypes. For instance, orthotopic transplantation in immunocompetent mice identified SOX17 as orchestrator of immune-evasive programs enabling tumor initiation^49^. Similarly, a stepwise melanoma model revealed that mutational combinations shape the cellular composition of the microenvironment and transcriptional states of individual cell types^50^. Notably, KAT organoids exhibited transcriptional features overlapping with those observed in a subset of IFN-low ATK CRCs, which are associated with poor responses to ICIs. While mutation order cannot be inferred from these clinical samples, these observations suggest a potential link between mutation-order-associated transcriptional states and tumor immune phenotypes. Further investigation in larger and clinically annotated cohorts will be important to determine the broader clinical relevance of these findings. Together, these results implicate the order of mutation acquisition in defining clinical outcomes.

Although our CRISPR-based murine KAT organoids closely resemble the IFN-low AKT human CRCs, our system still have some limitations in fully recapitulating human CRC tumorigenesis. First, due to the technical constrains, our CRC tumorigenesis model initiates with *Kras* mutation, which, although indeed occur early in human CRC, is rarely the initiating event in human CRC tumorigenesis^51,52^. Thus, our findings are contextualized with the *Kras*-first framework. Importantly, recent evidence suggests that oncogenic *Kras* activation can reshape the selective landscape for subsequent driver events, including *Apc* alterations. Consistent with this notion, our findings indicate that *Kras^G12D^* activation may establish a permissive cellular state that influences the phenotypic consequences of subsequent mutations. Future studies examining *Apc/Trp53* mutation ordering in *Kras* wild-type backgrounds will be important to define the broader generalizability of these findings.

Additionally, due to the time-consuming and laborious nature of establishing isogenic clones, we employed only two isogenic clones per genotype in all functional assays. Nevertheless, the consistent phenotypes observed in these experiments provide robust evidence for our key findings. Future work would benefit from the analysis of additional clones to further reinforce generalizability. Secondly, while CRC progression involves additional drivers (e.g., *SMAD4, PIK3CA*) and passenger mutations that collectively influence early transformation^11,53,54^, this study focused exclusively on altering the order of two key drivers (*Apc* and *Trp53*) during early carcinogenesis. This may not fully capture the broader complexity of CRC development. Third, short-term *in vitro* mutagenesis and subcutaneous transplantation models may not fully recapitulate the long-term evolutionary processes and tissue-specific microenvironmental interactions during human CRC development, where genetic, epigenetic, and microenvironmental changes cooperatively reshape clonal dynamics^33,43^. Although subcutaneous transplantation provides a tractable platform for assessing tumorigenic potential under controlled immune conditions, the cutaneous immune microenvironment differs substantially from that of the intestinal mucosa. Future studies incorporating orthotopic transplantation and systematic profiling of microenvironmental remodeling will help further clarify how mutation order shapes early CRC evolution.

In summary, our work systematically dissected the essential genetic changes during early colorectal carcinogenesis, establishing relationships between specific mutation combinations, their order, and the resulting malignant phenotypes as well as transcriptomic features. This study advances organoid-based modeling of multistep tumorigenesis, provides new insights into the origins of tumoral heterogeneity, and establishes an experimental framework for further mechanistic and clinically relevant studies.

## Acknowledgements

This work was supported by the National Key R&D Program of China (Grant No. 2022YFC3400400 to W.C.), National Natural Science Foundation of China (Grant No. 32470590 to L.F.), Shenzhen Key Laboratory of Gene Regulation and Systems Biology (Grant No. ZDSYS20200811144002008 to W.C.), Shenzhen Science and Technology Program (Grant No. KQTD20180411143432337 to W.C.). We thank the members of the Chen lab for insightful discussions and suggestions during the process of this investigation. The authors acknowledge the Center for Computational Science and Engineering of SUSTech for the support on computational resources and acknowledge the Laboratory Animal Center of SUSTech and the SUSTech Core Research Facilities for technical support.

## Author contributions

W.C., Q.Z. and L.F. conceived the project and supervised the study. Y.L. performed the experiments with assistance from D.D., Z.S., Z.H. and Y.T., and X.X. performed the computational analyses with assistance from Y.L.. Q.Z., W.C., L.F., and Y.L. wrote the manuscript with input from all authors. X.X., D.D. and Z.S. reviewed and revised the manuscript.

## Material and Methods

### Reagent Table

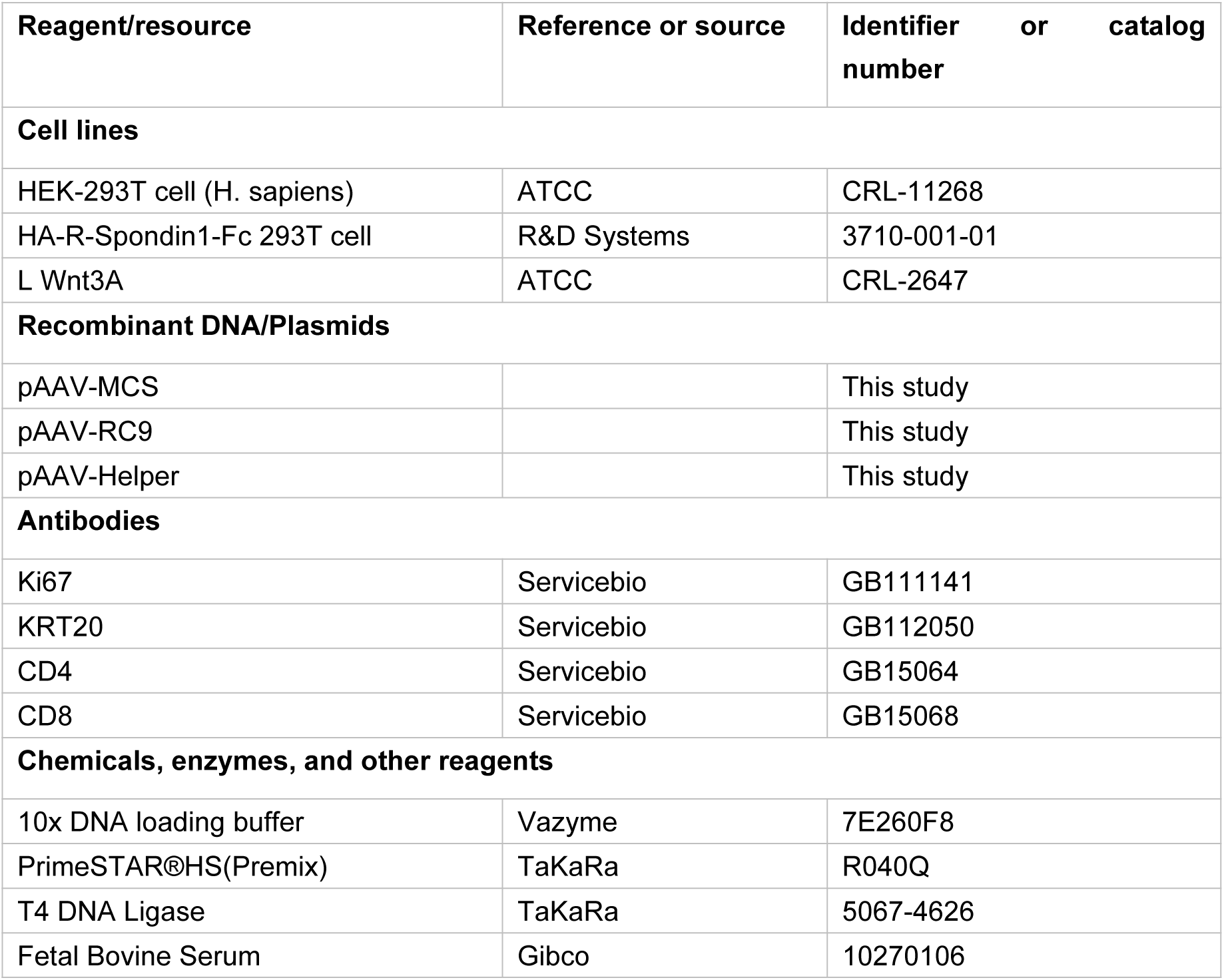
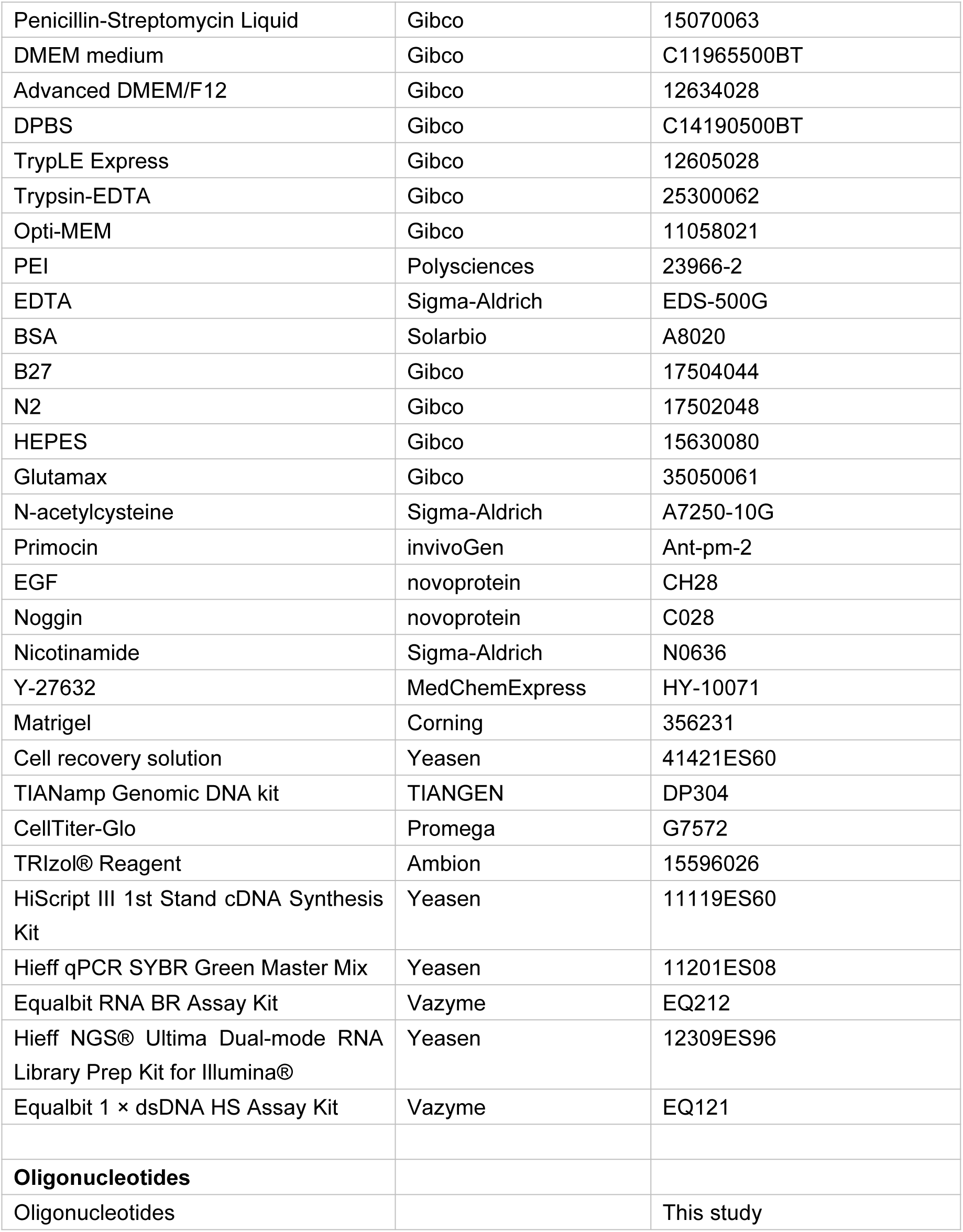

### Mice

Mice carrying a LoxP-Stop-LoxP (LSL) allele of *Kras^G12D^* (*Kras^LSL-G12D/+^*, NM-KI-190003) were bred with mice carrying homozygous Rosa26^CAG-LSL-Cas9-tdTomato/+^ (GemPharmatech, strain no. T002249) to generate mice carrying these two genotypes. Kras^LSL-G12D/+^; Rosa26^CAG-LSL-Cas9-tdTomato/+^ mice were generated after several generations of inbreeding. For *in vivo* tumor transplantation, immunodeficient B-NDG (NOD*-Prkdc^scid^IL2rg^tm^*^1^*/*Bcgen) and wild-type C57BL/6 mice were purchased from Biocytogen Co., Ltd.. All mice were housed in the specific pathogen-free animal care facility at thermal neutral temperature under a 12 h/12 h light/dark cycle. All animal work was performed in accordance with protocols approved by the Laboratory Animal Welfare and Ethics Committee at Southern University of Science and Technology (SUSTech, Shenzhen, China).

### Culture of mouse intestinal organoid

Primary mouse intestinal crypts were isolated and cultured in epidermal growth factor/Noggin/R-spondin1 (ENR) medium as described previously with modifications^29,55^. Briefly, the intestinal segment was dissected into 2-5 mm pieces and incubated in dissociation buffer (5 mM EDTA in DPBS supplemented with 1% Penicillin/Streptomycin) at 4 °C for 30 minutes with agitation. Next, the tissue pieces were removed from dissociation buffer and washed twice with 1x DPBS, and mechanically dissociated by pipetting in 0.1% BSA in 1x DPBS. After several rounds of dissociation, the isolated crypts were collected, strained and centrifuged, then resuspended in Matrigel (Corning, 356231) at an appropriate density for seeding, and ENR medium was added.

The culture medium ENR contains advanced DMEM/F12 medium (Gibco, 12634028), with supplement of B27 (Gibco, 17504044), N2 (Gibco, 17502048), Glutamax (Gibco, 35050061), Hepes (Gibco, 15630080), N-acetylcysteine (Sigma-Aldrich, A7250-10G), Primocin (invivoGen, Ant-pm-2), EGF (Novoprotein, CH28), Noggin (Novoprotein, C028) and R-spondin1 conditioned medium (20% v/v, produced using HA-R-Spondin1-Fc 293T cells, R&D Systems, 3710-001-01). For viral infection, organoids were cultured with medium ENRW, which was further supplemented with WNT conditioned media (50% v/v, produced using stably transfected L-Wnt3A cells, ATCC® CRL-2647™) and nicotinamide (Sigma-Aldrich, N0636).

Organoids were passaged every 5-7 days with medium replacement every 2-3 days. For passaging, organoids were released from the Matrigel, physically disrupted or digested with TrypLE Express (Gibco, 12605028) for about 5 minutes into small clumps of cells, washed with pre-cooled advanced DMEM/F12 and replated. Mycoplasma detection was regularly performed and resulted negative.

### Stable organoid line generation

Cre-mediated recombination was used to generate oncogenic activation of *Kras^G12D^*, *Cas9* and tdTomato expression, and CRISPR-Cas9 technology was then used for specific gene knockout. The adeno-associated virus serotype 9 (AAV9) packing system (pAAV-MCS, pAAV-RC and pHelper plasmids) was purchased from Strategene (La Jolla, CA, USA). A U6-filler-gRNA-scafold fragment together with a GFP/BFP expressing fragment were firstly cloned into pAAV-MCS between left and right ITR, then *Tp53*-gRNAs and *Apc*-gRNAs were cloned into these plasmids respectively to get gRNAs and fluorescent reporter constructs. The sgRNA sequence was determined by the CRISPR Design Tool (http://crispr.mit.edu/). An N6 random hexamer after reporters was also inserted to the plasmids to create a series of pAAV-gRNA-Reporter-Barcode constructs. All primer and gRNA sequences are listed in Table S3. All constructs were confirmed by Sanger sequencing. Reporter expression was confirmed in HEK-293T cells by transient transfection using polyethylenimine (PEI, Polysciences, 23966-2).

AAV9 was produced according to the manufacturer’s instructions. In detail, 120 µg of constructs (pHelper, pAAV-RC9, and pVector, equal molar ratio) mixed with 360 µg of PEI were required for transfection of HEK-293T cells in one 15-cm dish. Cells were collected 72 hours later and lysed by freeze-thaw cycles. Viruses were concentrated by PEG and extracted by chloroform. Extracted virus were then dissolved in HBS (50 mM HEPES, 150 mM NaCl, 25 mM EDTA pH 8.0) and titrated using quantitative PCR (qPCR) as previously described^56^. AAV-Cre was from Genomeditech (Shanghai, China). The virus was used in aliquots to avoid multiple freeze-thaw cycles.

For AAV infection, the organoids were dissociated into single cells or small cell clusters, and the cells were resuspended with viral solution in ENRW media and spread on 50% Matrigel-coated plates. After incubation overnight, living cells attached to Matrigel were collected to seed back into fresh Matrigel in dome shape. When cells grow out, the infected organoids were sorted by Fluorescence activated cell sorting (FACS) according to the reporter fluorescence expression and replated into a new drop of Matrigel for enrichment. Single clones were then picked from single cells derived organoids from positively infected organoids after FACS. Genomic DNAs of engineered organoids were extracted using TIANamp Genomic DNA kit (TIANGEN, DP304) and used for validation of targeted mutations by PCR amplification of targeted loci. PCR products were directly sequenced or cloned into a pMD19T cloning vector according to the manufacturer’s instructions (Takara, 6013), and mutation status of isogenic clones were validated according to the sequencing results.

### Organoid formation assay

Organoids were dissociated by vigorous resuspension in TrypLE Express for 10 minutes at 37 °C, and the dissociated single cells were collected, washed and counted, then resuspended in Matrigel, and ENR medium supplemented with 10 μM Rho kinase inhibitor Y-27632 (MedChemExpress, HY-10583) was added. Organoid formation efficiency was calculated by dividing the number of organoids formed by the total number of single cells plated after 5 days.

### Organoid growth detection

Organoids were split at a ratio of 1:3 to 1:10 (recorded) during passages. Specifically, one well of organoids were split and resuspended in Matrigel equivalent for indicated wells of organoids. Subsequent passages followed the same split paradigm. At each passage, the cell number was determined by CellTiter-Glo (Promega, G7572) to generate a growth curve, and cell numbers during 4 passages were determined.

### *In vivo* transplantation and tumor dissection

The organoid lines were expanded and dissociated with TrypLE Express for 5-10 minutes at 37 °C. After dissociation, 5x10^5^ cells (experiment 1, injection of KA, KT, KAT and KTA organoids for comparison of tumor features) and 1x10^6^ cells (experiment 2, injection of KAT and KTA organoids for further comparison of tumor growth) were resuspended in 50% Matrigel diluted in ENR and injected subcutaneously into immunodeficient B-NOD (NSG; NOD-*Prkdc^scid^IL2rg^tm^*^1^) mice. For subcutaneous transplantation in immunocompetent mice, 1x10^6^ cells of KAT and KTA organoids were injected into C57BL/6 wild-type mice. Two independent experiments were performed, of which the second experiment here was conducted together with the second tumorigenesis experiment in immunodeficient mice mentioned above.

Tumor progression was documented three times every week from the time of inoculation to the experimental endpoint. Tumor dimensions were measured using calipers and tumor volume was calculated as 0.5 × length × width × width. Tumor outgrowth was defined as the successful formation of a palpable subcutaneous mass that persisted and showed progressive growth for at least two consecutive measurements post-injection. The latency to outgrowth was recorded as the time point when the tumor first became palpable and met these criteria. Animals were humanely euthanized according to the following criteria: clinical signs of persistent distress or pain, significant body-weight loss (>20%), tumor size exceeding 1,000 mm^3^, or the occurrence of tumor ulceration or failure of tumor formation during the experimental period. At the experimental endpoint, animals were sacrificed, and tumors were carefully collected, minced using surgical scissors. For the histological study, tumors were immediately fixed in 4% PFA until paraffin embedding. The samples were sectioned into 5-μm sections.

### RNA isolation and quantitative RT-PCR

Total RNA from organoids was extracted by RNA isolater Total RNA Extraction Reagent (Vazyme, R401). To generate cDNA, equal amounts of total RNA from organoids were incubated with cDNA Synthesis Master Mix (Vazyme, R223) according to the manufacturer’s instructions. cDNA was used for quantitative real-time PCR using 2x SYBR Master Mix (Vazyme, Q711). Gene expression was normalized to β-Actin RNA. Primer sequences used for gene expression analysis by qPCR are listed in Table S3.

### RNA library preparation

RNAs from indicated organoids or tumor cells were converted into cDNA libraries using the Hieff NGS® Ultima Dual-mode mRNA Library Prep Kit (Yeasen, 12309ES24). High-throughput sequencing was performed using Illumina NovaSeq 6000.

### Histology and immunohistochemistry

For histological analysis of organoids, Matrigel embedded organoids were collected using Cell recovery solution (Yeasen, 41421ES60) to remove the Matrigel and washed once with 1x DPBS with 0.1% BSA carefully, then fixed with 10% NBF for 30 min at RT and stored at 4 °C until paraffin embedding. The samples were sectioned in 5-μm sections.

For H&E staining of both organoids and tumor samples, sections were stained following the instructions of H&E staining Kit (Servicebio, G1212). For immunohistochemistry, antigen retrieval was performed using citrate buffer (Servicebio, G1218). Block solution (Servicebio, G2010) was used. Antibodies and respective dilutions used for immunohistochemistry are as follows: anti-KRT20 (1:3000, Servicebio, GB112050), rabbit polyclonal anti-Ki67 (1:8000, Proteintech, 27309-1-AP), anti-CD4 (1:600, Servicebio, GB15064), anti-CD8 (1:800, Servicebio, GB15068), and HRP conjugated secondary anti-rabbit (1:500, Servicebio, GB23303) or anti-mouse antibodies (1:500, Servicebio, GB23301). DAB substrate kit for visualization (Servicebio, G1212) was used. All stained sections were imaged using a TissueFAXS system (TissueGnostics, Vienna, Austria).

The proportion of KRT20-positive cells was quantified from immunohistochemistry (IHC) images using ImageJ Fiji software. Briefly, for each image of a complete tumor section, the total area of tumor cells per field was determined by total size of the tissue within the whole image. KRT20-positive cells were identified based on a DAB intensity threshold that was set to clearly distinguish specific signal from background, which was applied consistently across all compared groups. The percentage of KRT20-positive cells was calculated as (Area of KRT20-positive cells/Total Area of tumor tissue) × 100% for each field of view, and the results were presented as mean ± SEM across all samples.

### RNA-seq analysis

Raw reads were first processed using Cutadapt v 4.4 with parameters -m 20 -q 30 to trim adapter sequences^57^. Subsequently, the clean reads were aligned by STAR v 2.7.10b against the mouse reference genome (GRCm38) using Ensembl v99 annotations excluding pseudogenes in advance^58^. Unique mapped read counts per gene were determined using featureCounts v 2.0.1 with parameter -p -B -C -t exon^59^.

Genes with TPM > 1 in at least one condition were retained for further analysis, including PCA, differential expression analysis, and GSEA. Differential expression analysis between pairs of conditions was performed using DESeq2 with default settings^60^. Genes with an absolute Log2 Fold Change > 1 and an adjusted p-value (padj) < 0.05 were defined as DEGs, except for specially marked samples. MA-plots were generated using the R package ggpubr (https://rpkgs.datanovia.com/ggpubr/).

The list of DEGs was subjected to functional annotation by GO, KEGG, and GSEA with the mouse MSigDB hallmark dataset using R package clusterProfiler^61^. Pathway visualizations were created using the ggplot2 (https://ggplot2.tidyverse.org) package to generate dot plots.

Subsequent heatmap analyses were computed with genes that were significantly differentially expressed between conditions K, KA, KT, KAT, and KTA versus SKC. Normalized counts were log2-transformed and row z-score normalized prior to heatmap generation. To generate the summary gene expression dynamics shown in Figure 4F, the median normalized counts per gene set were used prior to log transformation.

### ssGSEA of clinical samples

The gene expression data for the CRC cohort were obtained from the sequencing data by laser-capture microdissection, which was deposited in Zenodo with accession number: 10927005. For single-sample GSEA (ssGSEA), CRCs with *APC*, *TP53*, and *KRAS* triple mutations were selected from a total of 58 CRC patients. Tumor intrinsic gene signatures representing CRC biology and the interaction with the TME were referred from the literature. Per-sample GSEA score for the KAT vs. KTA upregulated IFN signatures (designated as organoid hallmark geneset) were evaluated using ‘ssgsea’ method in GSVA^62^. Generation of box plots showing the distribution of NES was performed using ggplot2.

### Statistical analysis

Unless otherwise specified in the figure legends or Methods, all experiments reported in this study were repeated at least three independent times. All sample number (n) of biological replicates and technical replicates can be found in the figure legends. The images for H&E staining and immunohistochemistry represent one of ≥ 2 biological replicates unless otherwise stated. All values were presented as mean ± SEM. unless otherwise stated. Intergroup comparisons were performed using two-tailed unpaired t-tests or One-way analysis of variance (ANOVA) with post-hoc Tukey’s multiple comparison. P values or adjusted P values of < 0.05 were considered to be significant. Statistical analysis was performed by GraphPad Prism v10.4.2. Studies were not conducted blind except for all histological analyses. Statistical details are found in the figure legends.

## Data availability

The raw and processed bulk mRNA-seq data generated in this study have been deposited at the Gene Expression Omnibus (GEO) public repository under accession number GSE300119. The code used for data processing and analysis is available upon request, and the software used is listed in the Method.

---

**Supplementary Figure 1.**
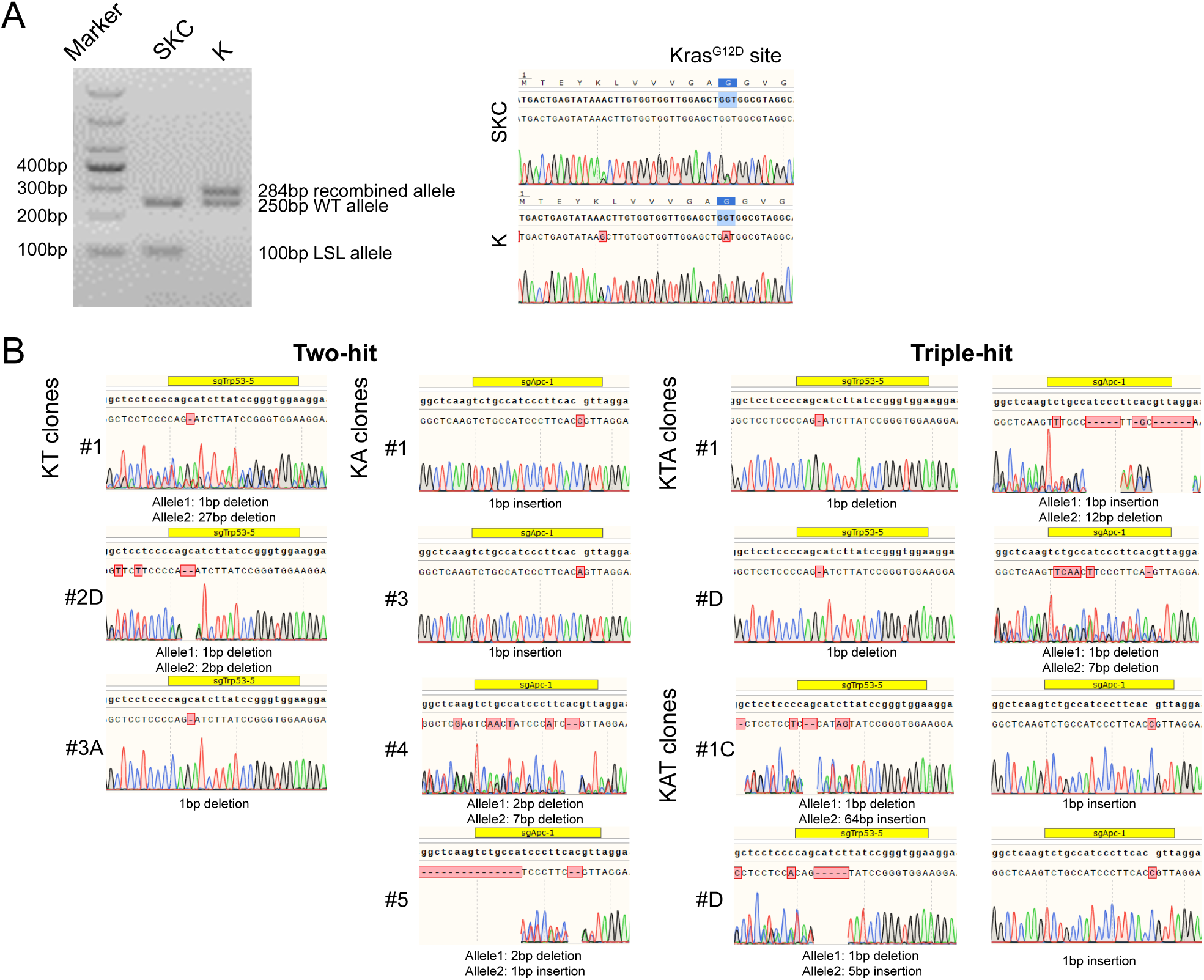
CRISPR-Cas9-mediated introduction of CRC mutations in murine intestinal organoids, related to Figure 1. A. Validation of excision of Stop elements and *Kras^G12D^* mutation on K organoids by PCR reaction targeting LoxP-Stop-LoxP site and *Kras* gene, and PCR products were used for electrophoresis (left panel) and Sanger DNA sequencing (right panel). Oncogenic GGT>GAT mutation is indicated in blue. B. Sequence analysis of the targeted *Trp53* and *Apc* exon in engineered organoids. Specific indels at expected locations were analyzed using Synthego’s Inference of CRISPR Edits (ICE) software, which were indicated under the alignment figures.

**Supplementary Figure 2.**
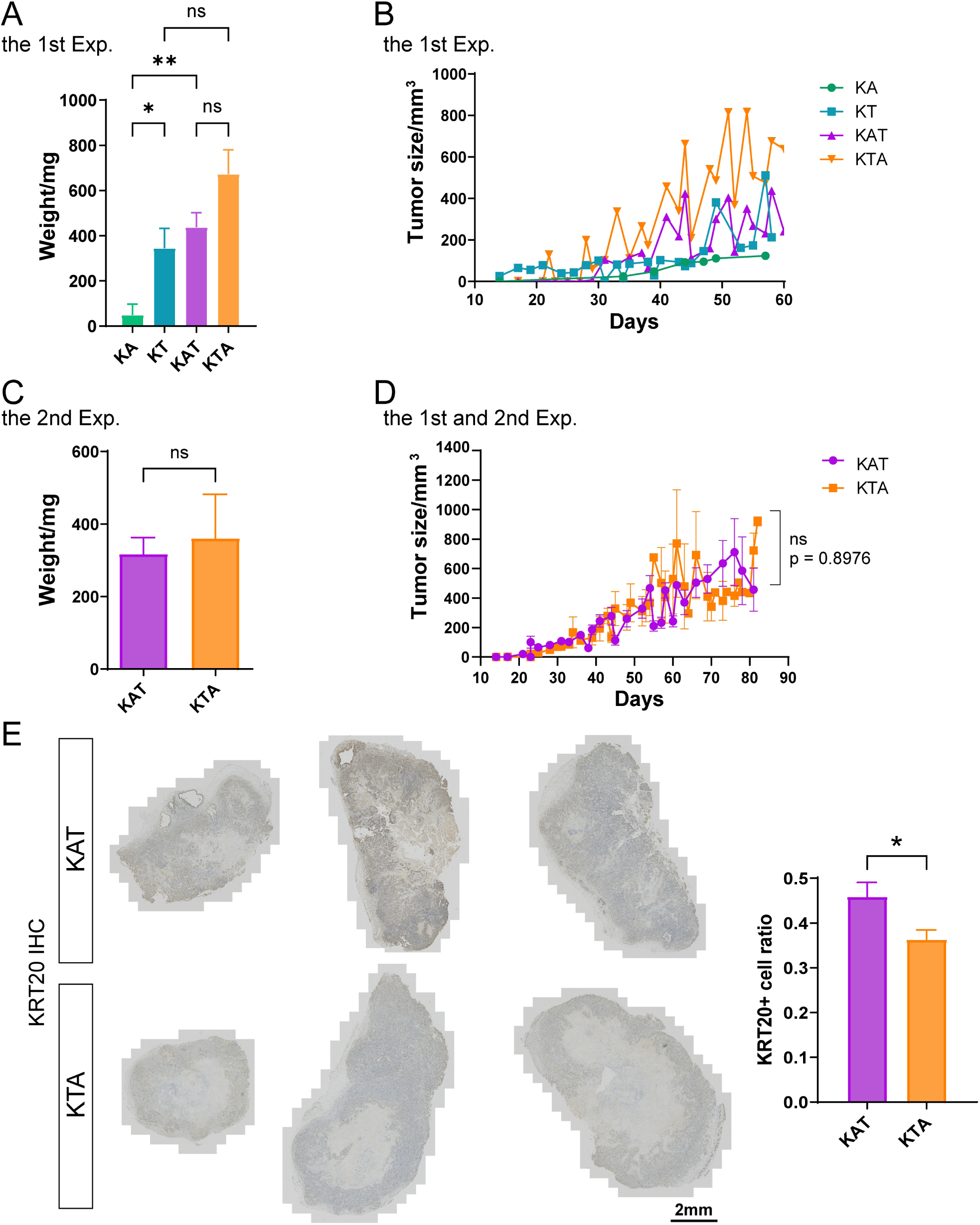
Mutation burden inversely correlated to tumor development kinetics and tumor size, related to Figure 3. A. Tumor weights from KA, KT, KAT and KTA organoids harvested at differed time-points (data from the first experiment). The analysis was performed with n = 6-9 injections for each genotype. B. Growth kinetics of KA, KT, KAT and KTA tumors overtime (data from the first experiment). The analysis was performed with n = 5 lesions for each genotype. C. Tumor weights from KAT and KTA organoids harvested at the same time-point (data from the second experiment). The analysis was performed with n = 6 tumors for KAT and KTA. D. Growth kinetics of KAT and KTA tumors from both Experiment 1 and 2 overtime. The analysis was performed with N = 12 tumors for KAT and N = 13 tumors for KTA. E. Representative images of immunostaining of Keratin 20 (KRT20) on KAT and KTA tumors, and right panel shows the KRT20 positive ratio in KAT and KTA tumors. Scale bar: 2mm. The analysis was performed with n = 6 tumors for KAT and KTA. Statistical analysis of A was done by One-way ANOVA, P values were indicated in the figure. Statistical analysis of D was done by Two-way ANOVA, adjusted P values: ns, P > 0.05. Statistical analysis of C and E was done by unpaired Students’ t-test, adjusted P values: ns, non-significant, P > 0.05; *, P < 0.05. The clones used in these analyses were listed in Table S5.

**Supplementary Figure 3.**
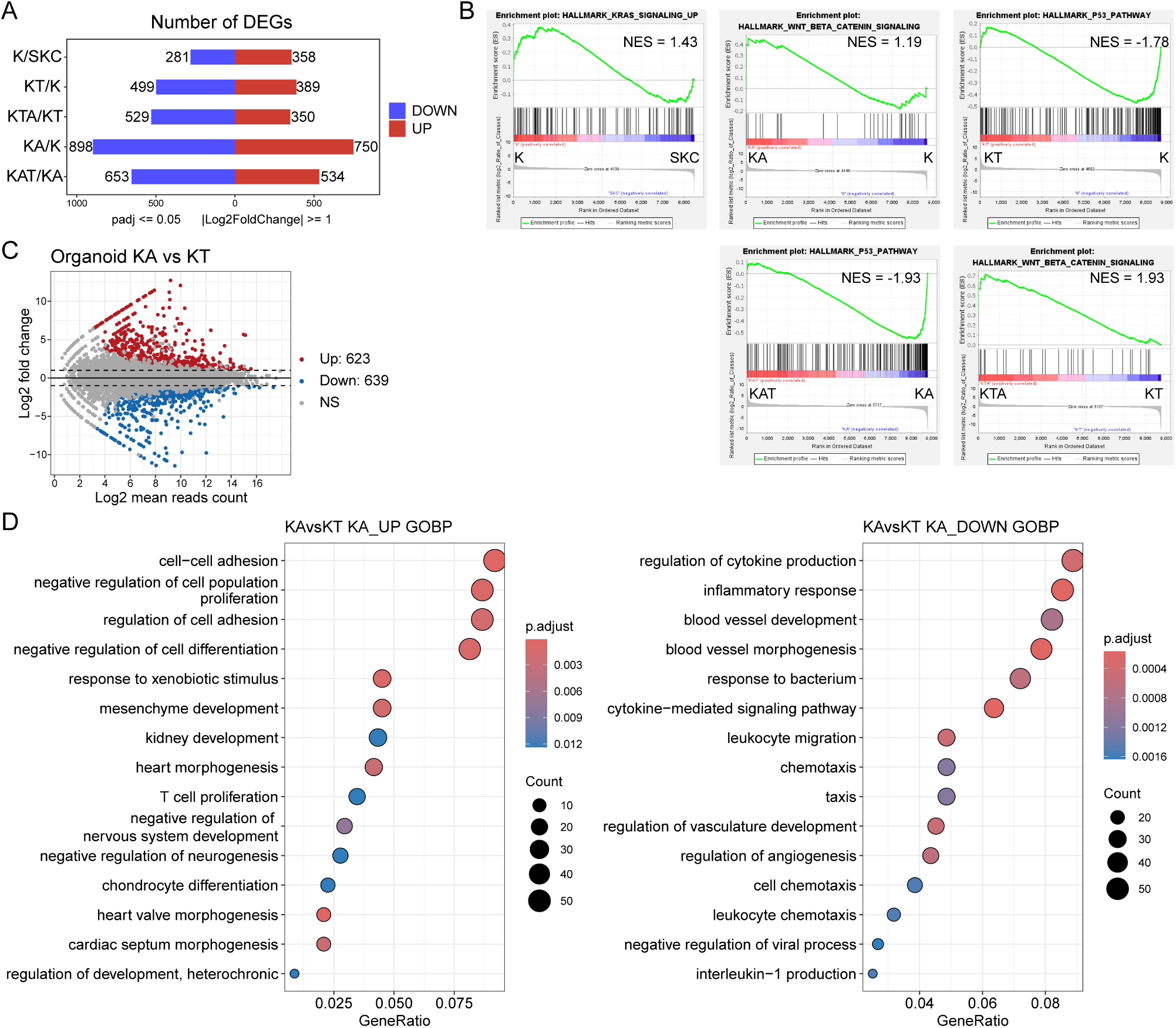
Stage-specific pathway alterations during stepwise mutagenesis and comparative analysis of KA and KT organoids, related to Figure 4. A. Bar plot showing the number of differentially expressed genes (DEGs) along each step of mutagenesis. The DEGs were defined as genes with LFC >= 1, padj <= 0.05. Red bars represent up-regulated genes, and blue bars represent down-regulated genes. B. GSEA results reflecting the transcriptional changes in core biological pathways upon *Kras^G12D^* activation and *Apc* or *Trp53* loss during each step of mutagenesis, with normalized enrichment score (NES) indicated at top right corner. C. MA-plot showing the DEGs between KA and KT organoids, with 623 up-regulated genes and 639 down-regulated genes. D. GO analysis showing the transcriptional differences between KA and KT organoids. The clones used in transcriptional analyses were listed in Table S5.

**Supplementary Figure 4.**
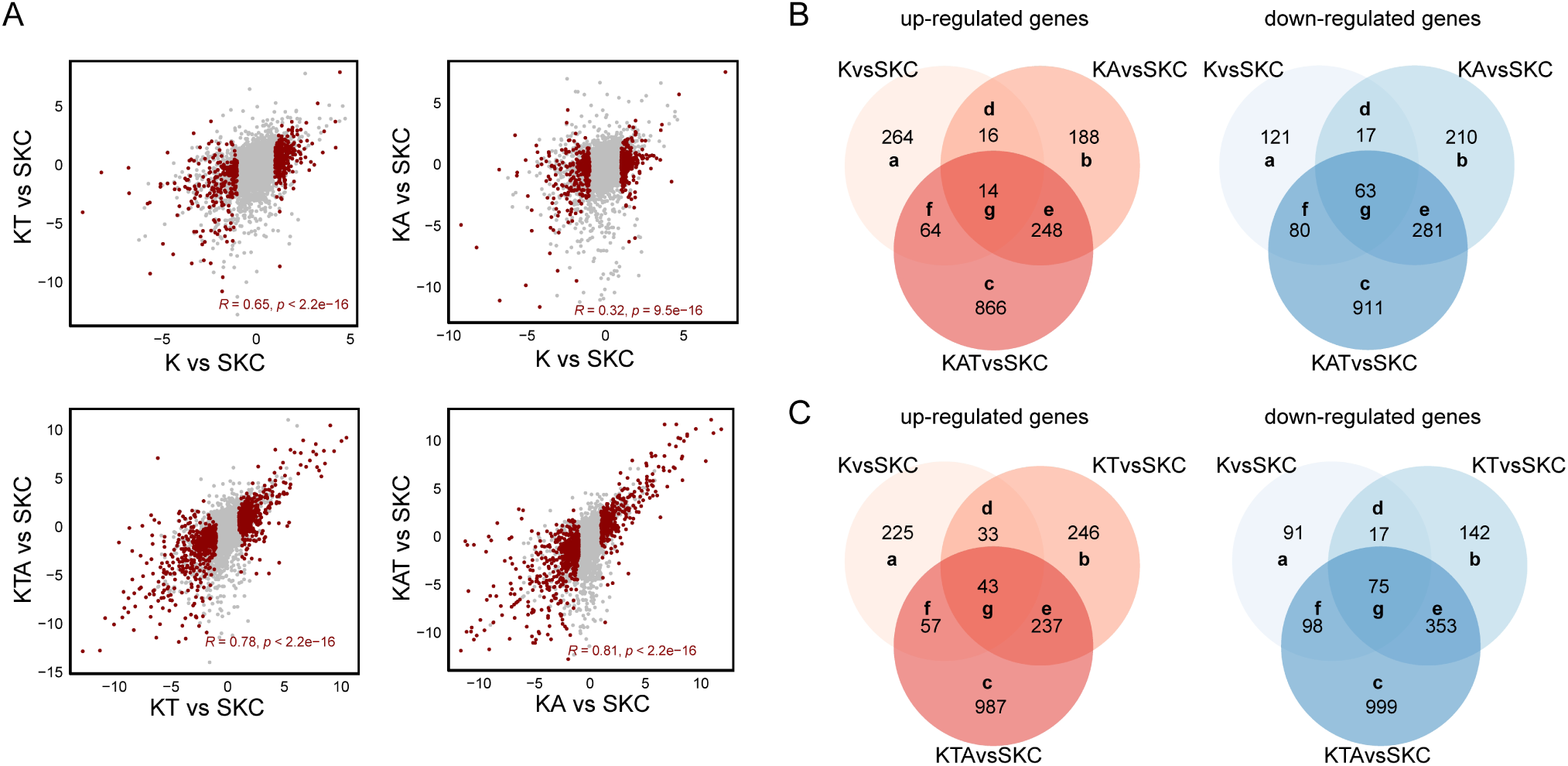
Transcriptional changes during mutation accumulation, related to Figure 4. A. Scatterplots of gene expression changes (LFC) showing correlation between the two comparisons indicated in the x and y axes of each plot. Significant changes (padj < 0.05) in the x axis comparisons are highlighted in red. The R values represent Spearman correlation values calculated from the LFC of genes with significant changes in the x axis comparison. B and C. Venn plots showing the up- (left panel) and down-regulated (right panel) genes across mutagenesis paths SKC>K>KA>KAT (B) and SKC>K>KT>KTA (C). The clones used in transcriptional analyses were listed in Table S5.

**Supplementary Figure 5.**
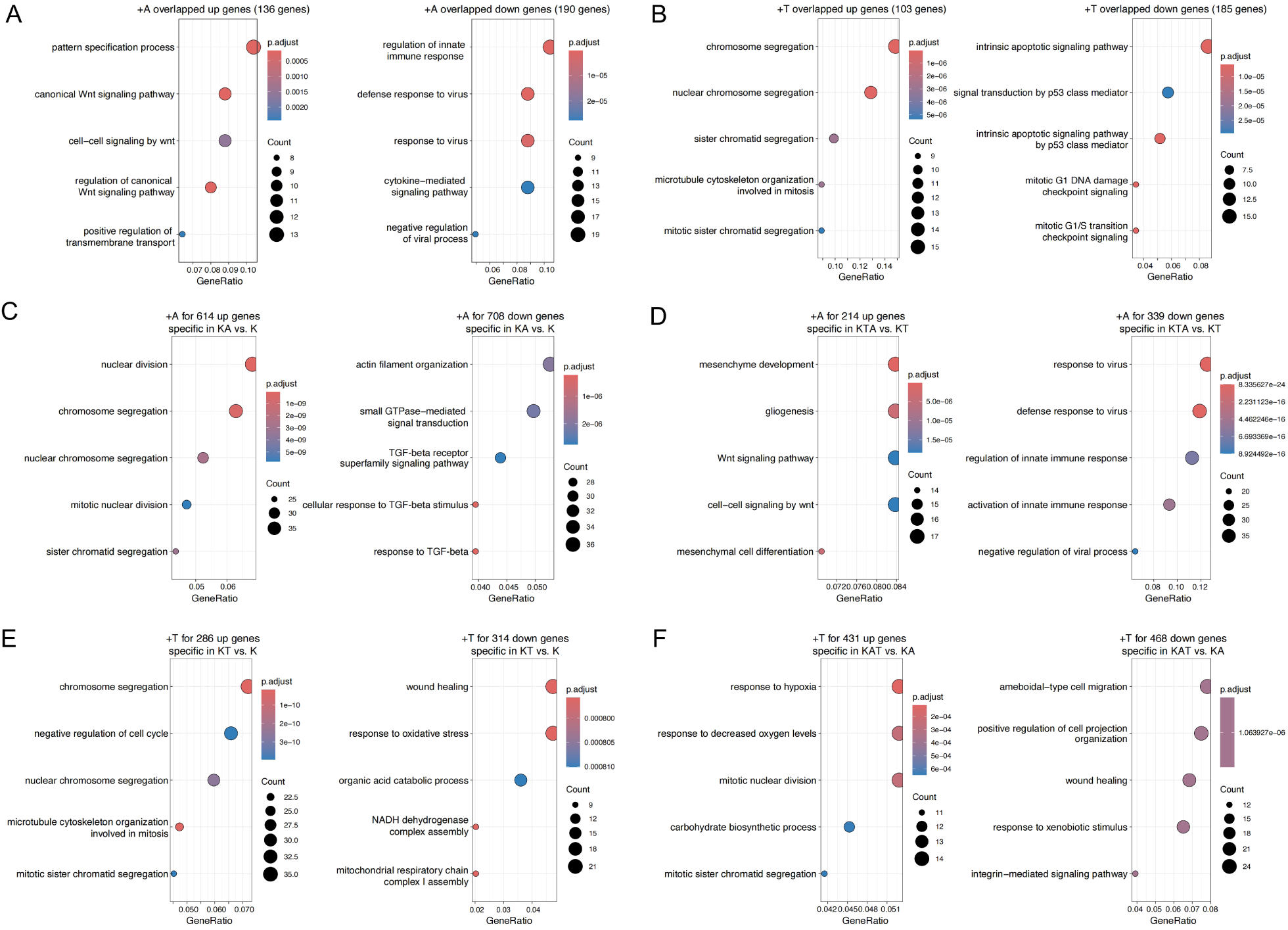
GO analysis of genes associated with conserved and context-specific transcriptional changes upon the same genetic mutation, related to Figure 5. A and B. GO analysis of overlapped up- and down-regulated genes at *Apc* inactivation (A) or *Trp53* deficiency (B). C and D. GO analysis of DEGs specifically altered at early *Apc* loss (K→KA) and late *Apc* loss (KT→KTA). E and F. GO analysis of DEGs specifically altered at early *Trp53* loss (K→KT) or late *Trp53* loss (KA→KAT). For all GO analysis results, top 5 biological pathways are shown. The clones used in transcriptional analyses were listed in Table S5.

